# Plasma membrane association and resistosome formation of plant helper immune receptors

**DOI:** 10.1101/2023.01.23.525201

**Authors:** Zaiqing Wang, Xiaoxiao Liu, Jie Yu, Shuining Yin, Wenjuan Cai, Nak Hyun Kim, Farid El Kasmi, Jeffery L. Dangl, Li Wan

## Abstract

Intracellular plant immune receptors, termed NLRs, respond to pathogen effectors delivered into plant cells. Activation of NLRs typically confers immunity. Sensor NLRs, involved in effector recognition, are either TIR-NLRs (TNLs) or CC-NLRs (CNLs). Helper NLRs, required for sensor NLR signaling, include CC^R^-NLRs (RNLs) and a special class of CNLs known as NRCs. Activated TNLs produce small molecules that induce an association between the EDS1/SAG101 heterodimer and the NRG1s helper RNLs. Auto active NRG1s oligomerize and form calcium signaling channels largely localized at the plasma membrane (PM). The molecular mechanisms of helper NLR PM association and effector induced NRG1 oligomerization are not well characterized. We find that both RNLs and NRCs require positively charged residues in the second and fourth helices of their CC^R^ or CC domain for phospholipid binding and PM association before and after activation, despite conformational changes that accompany activation. We demonstrate that effector activation of TNLs induces NRG1 oligomerization at the PM and that the cytoplasmic pool of EDS1/SAG101 is critical for cell death function. EDS1/SAG101 cannot be detected in the oligomerized NRG1 resistosome, suggesting that additional unknown triggers might be required to induce the dissociation of EDS1/SAG101 from the previously described NRG1/EDS1/SAG101 heterotrimer before subsequent NRG1 oligomerization, or that the conformational changes resulting from NRG1 oligomerization abrogate the interface for EDS1/SAG101 association. Our data provide new observations regarding dynamic PM association during helper NLR activation and underpin an updated model for effector induced NRG1 resistosome formation.

## Introduction

Plants employ both cell surface and intracellular immune receptors to detect pathogens. Cell surface Pattern Recognition Receptors (PRRs) recognize Pathogen-Associated Molecular Patterns (PAMPs) and initiate PAMP-triggered immunity (PTI) (1). Intracellular immune receptors known as Nucleotide-binding Leucine-rich repeat Receptor (NLR) proteins recognize corresponding effectors secreted by pathogens into plant cells and initiate effector-triggered immunity (ETI) (2). Plant NLRs express three main types of N-terminal domains: either a Toll/interleukin-1 receptor (TIR) domain, a coiled-coil (CC) domain, or an RPW8 (Resistance to Powdery Mildew 8)-like CC (CC^R^) domain. CC and CC^R^ domains adopt a conserved structure fold of 4 helical bundle (4HB) with the CC^R^ expressing an extra N-terminal region (3). TIR-NLR, CC-NLR and CC^R^-NLR are known as TNL, CNL and RNL, respectively (4). TNL and CNL recognize pathogen effectors to trigger cell death and immune responses and are thus termed sensor NLRs. Downstream of sensor NLRs, many seed plants also deploy helper NLRs to transduce signals. There are three described classes of helper NLRs. The Activated Disease Resistance 1 (ADR1) family, and the N Required Gene 1 (NRG1) family are highly conserved across the land plant phylogeny (5-7). ADR1s and NRG1s belong to the RNL family and are partially redundant downstream immune mediators of TNLs and some CNLs (8). NRCs are a greatly expanded helper NLR class in several plant families, including the Solanaceae (9). NRCs contain canonical CC domains at their N-termini. In *Nicotiana benthmiana* (*Nb*), three NRCs function redundantly downstream of diverse sensor CNLs to regulate cell death and immune responses (10).

Upon effector recognition, many tested sensor CNLs and TNLs oligomerize and undergo radical conformational changes. For example, the Arabidopsis CNL AtZAR1 and wheat CNL Sr35 form pentameric resistosomes and function as Ca^2+^ permeable channels to directly induce cell death and defense responses (11-15). In contrast, activated sensor TNLs form tetrameric resistosomes that are holoenzymes cleaving NAD^+^ to produce small signaling molecules (16-22). These small molecules bind to the heterodimers of the Enhanced Disease Susceptibility 1 (EDS1) with either Senescence-Associated Gene 101 (SAG101), or Phytoalexin Deficient 4 (PAD4) (19, 20). In Arabidopsis, AtEDS1 and AtSAG101 form a stable heterodimer and cooperate with the AtNRG1s to mainly control cell death, while AtEDS1 and AtPAD4 physically associate and function with the AtADR1s to mainly mediate bacterial growth restriction, resistance and transcriptional reprogramming (6, 7, 19, 20, 23, 24), though there are some overlapping functions (8). The auto-active AtNRG1 mutant and AtADR1 were demonstrated to oligomerize and function as Ca^2+^ permeable channel to induce cell death independent of the lipase-like proteins (3). It is currently unknown whether TNL activation is sufficient to trigger RNL oligomerization and whether EDS1/SAG101 and EDS1/PAD4 are present in NRG1 and ADR1 resistosomes, respectively.

Sensor NLRs are localized in their resting states to various places in the cell, likely in order to engage with the relevant appropriately localized effectors. Consistent with their proposed Ca^2+^ channel activities, some activated CNLs and RNLs exhibit PM localization (3, 12-15, 25, 26). The Arabidopsis CNL RPM1 associates with the PM and remains at the PM after its activation by PM targeted effectors that modify its guardee protein RIN4 (25, 27). The Arabidopsis CNL RPS5 also localizes to the PM via a N-terminal acylation signal and functions at the PM (26). By contrast, AtZAR1 distributes mainly in the cytoplasm in the resting state (13). Upon activation, AtZAR1 moves to, oligomerizes at and presumably protrudes through the PM to form an active Ca^2+^ channel (11, 12, 28). Plant helper NLRs including ADR1s, NRG1s and NRCs contain no predicted acylation signal but still localize at the PM before activation (3, 29, 30). AtNRG1.1 localizes to the PM, partially to ER membranes and in the cytosol when transiently over-expressed in *Nb* (3, 31). Arabidopsis ADR1s mainly localizes at the PM in a phospholipid dependent manner, since depletion of phosphatidylinositol-4-phosphate (PI4P) from the PM by overexpression of the yeast phospholipid-phosphatase Sac1p led to a mis-localization of ADR1 and loss of cell death activities (29). During activation by *Phytophthora infestans* infection, NbNRC4 accumulates at the extrahaustorial membrane (EHM) (30). When activated by sensor NLRs, NbNRC4 forms puncta mainly at the EHM and, to a lesser extent, at the PM; a constitutively autoactive NbNRC4 mutant mainly exhibits PM-associated punctate distribution, similar to autoactivated AtNRG1.1 (3, 30). Given that AtNRG1.1 CC^R^ cell death activity is also affected when co-expressed with the yeast phospholipid-phosphatase Sac1p (29) and that NbNRCs do also not possess any N-terminal acylation site or predicted transmembrane anchor, it is very likely that NRG1s and NRCs also associate with PM in a phospholipid dependent manner. However, how exactly the three subtypes of helper NLRs interact with phospholipids is unknown.

Here we show that the three classes of helper NLRs directly interact with phospholipids via conserved positively charged residues in the second and fourth helices of their CC^R^ or CC domain to anchor at the PM in both resting and active states, despite undergoing conformational changes accompanying activation. Consistent with the previously reported activation mimic AtNRG1.1 D485V (hereafter, AtNRG1.1 DV) (3, 31), effector activation of an upstream TNL induces AtNRG1.1 oligomerization and puncta formation at the PM. Interestingly, a cytoplasmic AtEDS1/AtSAG101 fraction is important for the AtNRG1.1 cell death function, but this heterodimer cannot be detected in the oligomerized AtNRG1.1 resistosome. These results suggest that the AtEDS1/AtSAG101/AtNRG1.1 heterotrimer induced by TNL activation represents an intermediate state before dissociation of AtEDS1/AtSAG101 and subsequent oligomerization of AtNRG1.1 or that the conformational changes resulting from AtNRG1 oligomerization abrogate the interface for AtEDS1/AtSAG101 association.

## Results

### Positively charged residues in α4 of the 4HB contribute to NRG1 and NRC phospholipid binding and PM localization before and after activation

We previously demonstrated that the AtADR1 CC^R^ domain binds to phospholipids *in vitro* (29), but it remained unknown which residues in AtADR1 CC^R^ were responsible for that interaction. Assuming that the ADR1 and NRG1 CC^R^ domains and potentially the NRC CC could use the same mechanism for phospholipid binding and PM association, we performed sequence alignment with ADR1s, NRG1s and NRCs from Arabidopsis and *Nb* to identify potential conserved, positively charged residues that might bind negatively charged phospholipids. We found an absolutely conserved lysine residue corresponding to K100 in AtNRG1.1 (Figure S1A). This lysine residue is highly conserved in ADR1s, NRG1s and NRCs across different plant species (Figure 1A). We did not observe conservation of this amino acid in AtRPS5 which features an N-terminal acylation signal for PM localization, in AtRPM1 which relies largely on association with RIN4 for PM association or in AtZAR1 which mainly localizes in the cytoplasm when overexpressed in *Nb* (Figure S1A and S1B). This observation further suggests a specific and potentially functional role for the AtNRG1.1 K100 equivalent in PM-localization of helper NLRs. Lipid-strip binding assays showed that purified AtNRG1.1 CC^R^ (aa 1-124) protein specifically binds to phospholipids including PI4P, while K100E completely abolished phospholipid binding. We note that even 2 or 5 times more K100E protein could not restore the binding to AtNRG1.1 CC^R^ wide-type (WT) level (Figure 1B). K100E was still active in triggering cell death in *Nb*, though this phenotype was weaker than the control, AtNRG1.1 DV (Figure 1C and 1D). Considering that positively charged residues adjacent to K100 may also contribute to phospholipid binding and function, we mutated positively charged residues close to K100 in the α4 helix (Figure S1C). A quadruple mutant K100E/R103E/K106E/K110E (hereafter, g4m) and a penta mutant R99E/K100E/R103E/K106E/K110E (hereafter, g5m) completely suppressed the cell death activity of AtNRG1.1 DV (Figure 1C and 1D). The four single mutants including R99E, R103E, K106E and K110E showed obviously or slightly reduced binding towards phospholipids *in vitro* (Figure 1E). Blue-Native PAGE assays showed that K100E and g4m properly oligomerized in the context of AtNRG1.1 DV, while g5m oligomerization was hindered (Figure 1F, also showing equal mutant accumulation on SDS-PAGE immunoblot). Both g4m and g5m strongly affected PM locations of both AtNRG1.1 WT and DV as demonstrated in both Laser Confocal Microscopy (hereafter, confocal) and PM fractionation assays, while K100E had a slight effect (Figure 2A, 2B, 2C and 2D). Given that the cell death function of AtRNG1.1 DV affects protein accumulation, PM fractionation experiments on AtRNG1.1 DV were performed in the context of the previously identified loss-of function mutant ΔN16 (deletion of residues 2-16) (Figure 2D) (3). In confocal assays, DV K100E and DV g4m maintained the ability to form puncta at the PM, as observed for AtNRG1.1 DV, but not DV g5m (Figure 2C, S2 and 2E), consistent with their abilities to oligomerize in Blue-Native PAGE assays.

**Figure 1.**
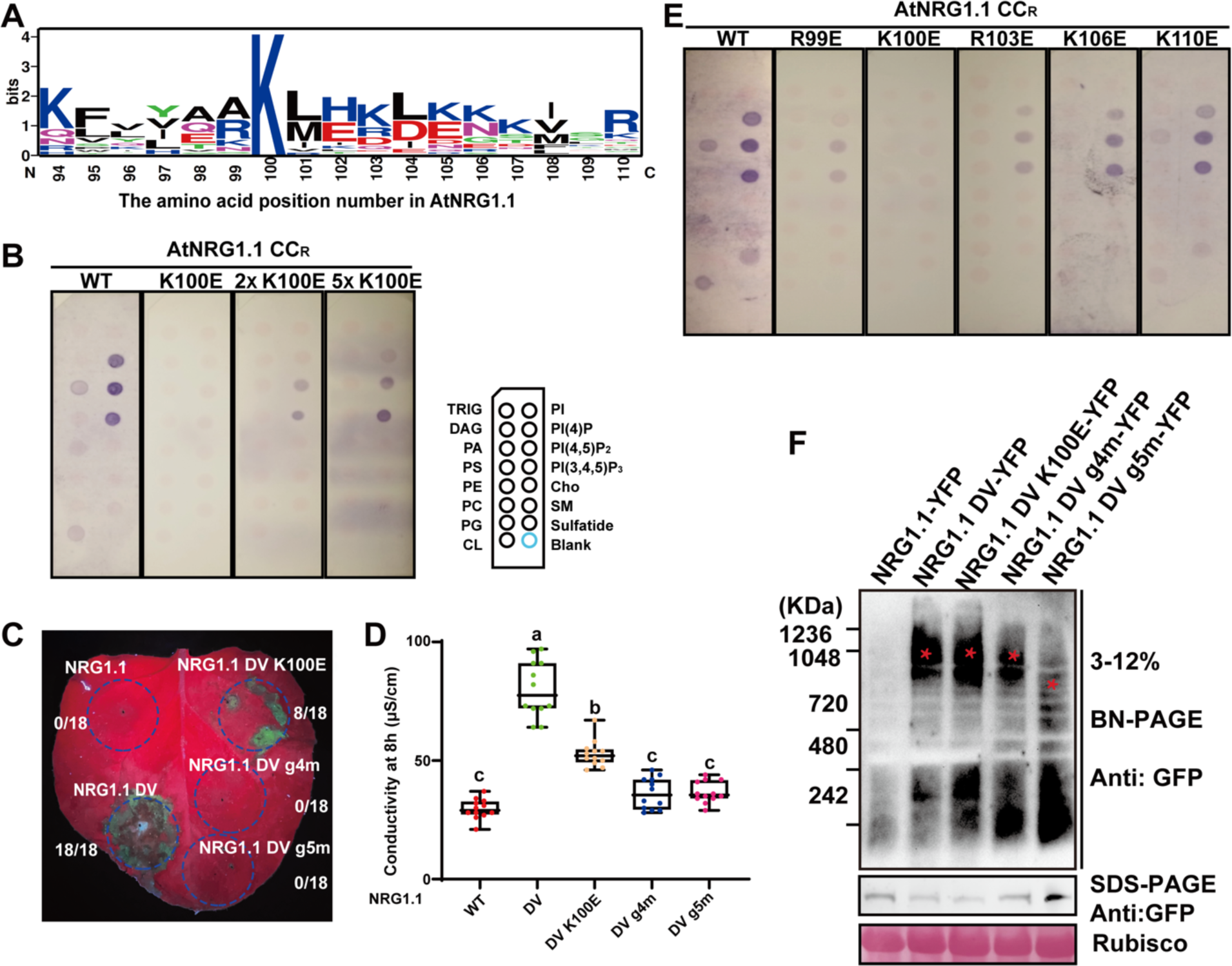
Positively charged residues in the α4 helix of 4HB affect AtNRG1.1 CC^R^ phospholipid binding and AtNRG1.1 DV function. **(A)** Sequence logos showing the conservation of AtNRG1.1 K100 in three helper NLR families from different plant species. The alignment was performed with ClustalX and sequence logos were generated using WEBLOGO. **(B)** Lipid strip binding assay using purified proteins of AtNRG1.1 CC^R^ (aa 1-124) and K100E. See *Methods* for lipid definitions. **(C)** Cell death phenotypes of AtNRG1.1 DV and mutants with altered PM localization in *Nb* at 32h post infiltration. (**D**) Quantification of ion leakage from the cell death phenotypes in (C). (**E**) Lipid strip binding assay using purified proteins of AtNRG1.1 CC^R^ 1-124 and mutants in the α4 helix. **(F)** BN-PAGE analyses of the oligomerization of AtNRG1.1 DV mutants with altered PM localization. Red asterisks indicate oligomerized AtNRG1.1 DV or mutant derivatives.

**Figure 2.**
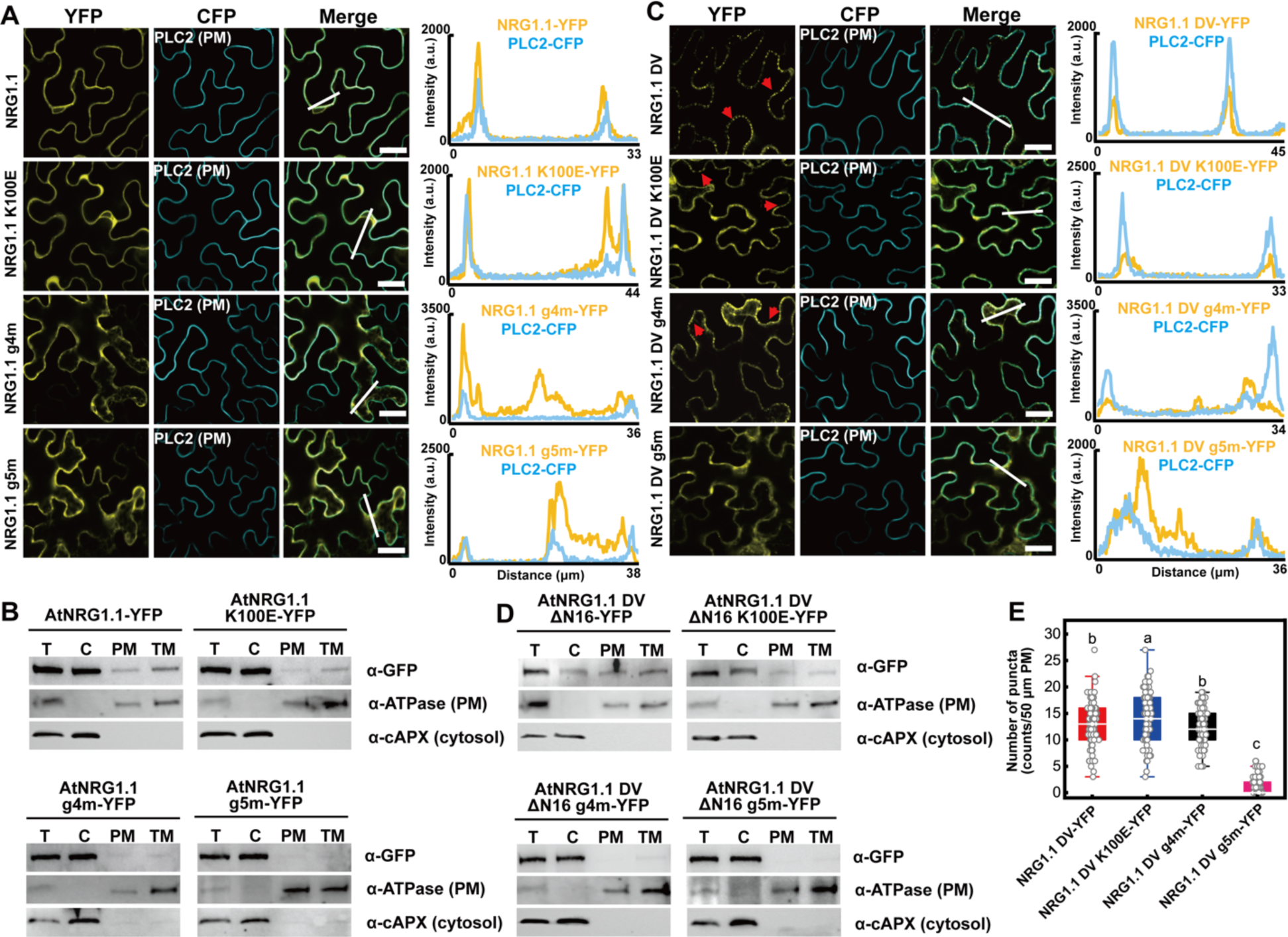
Positively charged residues in the α4 helix of 4HB affect AtNRG1.1 and DV PM association. Confocal microscopy analyses on the mutations of the positively charged residues affecting PM localization of AtNRG1.1 (**A**) and AtNRG1.1 DV (**C**). The indicated proteins fused with a C-terminal YFP were transiently co-expressed with the PM marker PLC2 fused to CFP in *Nb* leaves and confocal images were taken at 32-36h post infiltration. Confocal images are single plane secant views. Merge means merged image between YFP and CFP images. The red arrow heads indicated puncta. Fluorescence intensities were measured along the white line depicted in the merge images. Bars, 25 µm. Membrane fractionation assays on the positively charged residues affecting AtNRG1.1 (**B**) and AtNRG1.1 DV ΔN16 (**D**) protein accumulation at the PM. **(E)**, Quantification of puncta observed in (C).

The corresponding lysine residue is not conserved generally across CNLs, but is conserved in helper NRCs including the NRC negative regulator NbNRCX (Figure S1A) (32). The conserved lysine residue in the α4 helix of the Arabidopsis RNLs corresponds to K84 in NbNRC4. K84E/K87E/K89E/K91E/R94E (hereafter, c5m) abolished the phospholipid binding of NbNRC4 CC *in vitro* (Figure 3A) and suppressed the cell death activity of NbNRC4 D478V (hereafter, NbNRC4 DV) (Figure 3B, 3C and S3A). Moreover, c5m also attenuated the PM localization of both NbNRC4 WT (Figure 3D and 3E) and NbNRC4 DV (Figure 2F). Given the weak fluorescence signal of NbNRC4 DV due to cell death starting from 24h post infiltration, the confocal assay on NbNRC4DV was performed in the context of the previously identified loss-of-function mutant L9E (Figure 2F) (33). NbNRCX exhibited strong PM localization, while K84E/K89E/K92E/K93E (hereafter, x4m) exhibited reduced PM localization (Figure 2G and 2H). Hence, the positively charged residues in the α4 helix of the 4HB contribute to NRG1 and NRC phospholipid binding and PM localization before and after activation, suggesting a conserved mechanism.

**Figure 3.**
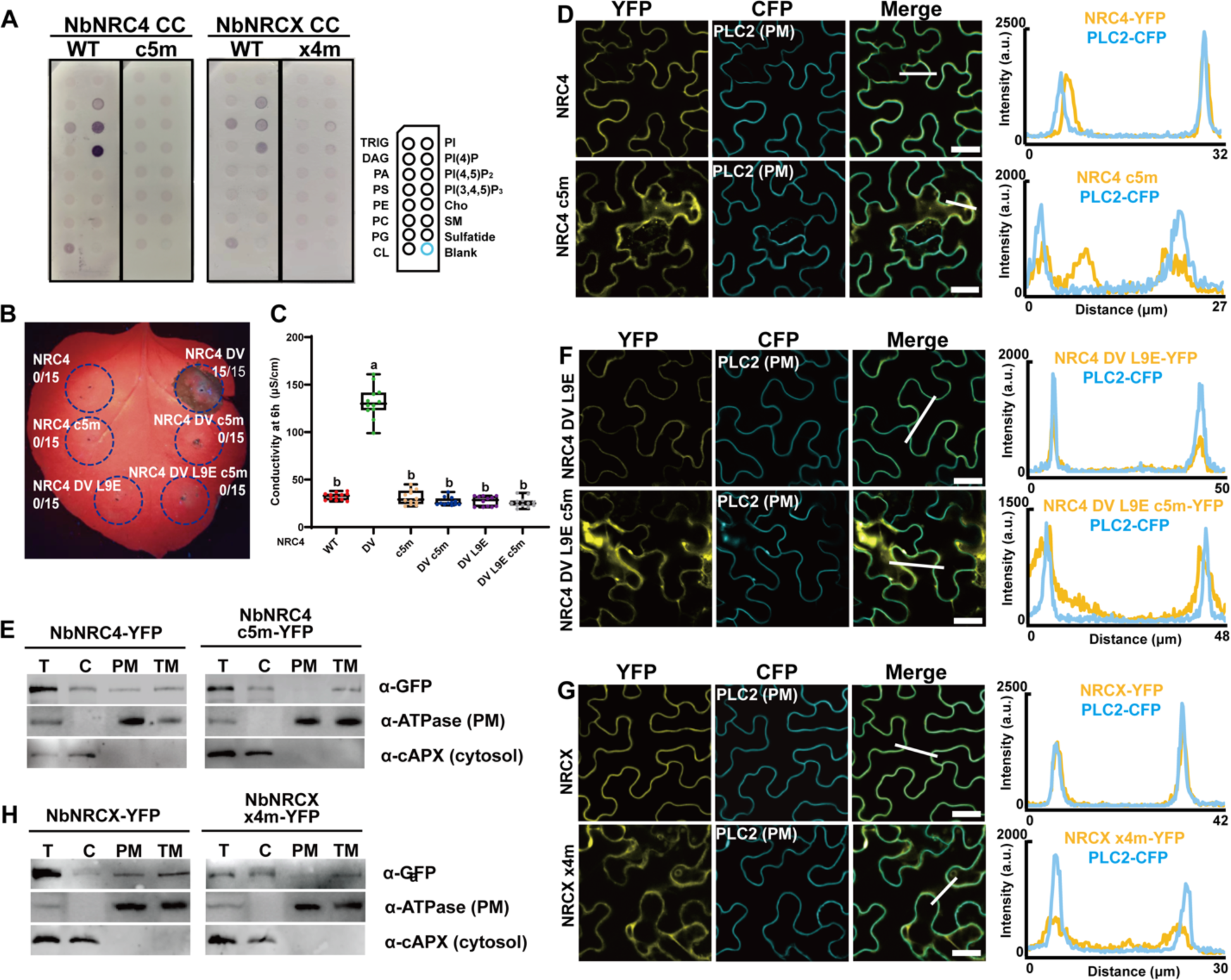
Mutations of the positively charged residues in the α4 helix of 4HB affect NbNRC phospholipid binding, PM localization and function. **(A)** Lipid strip binding assay using purified proteins of NbNRC4 CC (aa 1-127) and its c5m mutant and NbNRCX CC (aa 1-129) and its x4m mutant. See *Methods* for lipid definitions. **(B)** Cell death phenotypes of an NbNRC4 DV activation mimic and mutants with altered PM localization in *Nb* 32h post infiltration. (**C**) Quantification of ion leakage from the cell death phenotypes in (B). Confocal microscopy analyses on the mutations of the positively charged residues affecting PM localizations of NbNRC4 **(D)**, NbNRC4 DV **(F)**, and NbNRCX **(G)**, respectively. The indicated proteins fused with a C-terminal YFP were transiently co-expressed with the PM marker PLC2 fused to CFP in *Nb* leaves and confocal images were taken at 32-36h post infiltration. Images are single plane secant views. Merge means merged images between YFP and CFP images. Fluorescence intensities were measured along the white line depicted in the merge images. Bar, 25 µm. Membrane fractionation assays on the positively charged residues affecting NbNRC4 (**E**) and NbNRCX (**H**) protein accumulation at the PM.

### Positively charged residues in the α2 and α4 helices of 4HB collectively affect ADR1 PM localization before and after activation

Mutations in the α4 helix of the 4HB abolished the cell death function of AtNRG1.1 DV and NbNRC4 DV, but only partially affected PM localization, indicating that additional residues may also contribute to phospholipid binding. To further investigate if additional residues indeed contribute to phospholipid binding, we chose to focus on AtADR1-L1 because of its distinct PM localization compared to AtNRG1.1 and NbNRC4 (29). The conserved lysine residue in the α4 helix corresponds to K99 in AtADR1-L1. The quadruple mutant R98, K99, K102, K105 (hereafter, r4m) in α4 strongly attenuated the phospholipid binding of AtADR1-L1 CC^R^ *in vitro* (Figure 4A) and AtADR1-L1 D489V (hereafter, AtADR1-L1 DV) cell death activity (Figure 4B and 4C, with protein accumulation controls in S3B). r4m affected the PM localization of both AtADR1-L1 WT (Figure 4D and 4E) and AtADR1-L1 DV (Figure 4F). Further search for conserved positively charged residues in the 4HB of RNLs identified a highly conserved K or R residue corresponding to K30 in AtADR1-L1 and K35 in AtNRG1.1 (Figure S1A and 4G), and the corresponding positions in NRCs are also largely conserved (Figure S1A). A triple mutant of K30 and adjacent positively charged residues, R28E/K30E/K34E (hereafter, r3m), reduced the phospholipid binding of AtADR1-L1 CC^R^ *in vitro* (Figure 4A) and suppressed the cell death phenotype of AtADR1-L1 DV (Figure 4B and 4C), suggesting a critical role of the positively charged residues in the α2 helix. Confocal and membrane fractionation assays further demonstrated that r3m affected the PM localization of both AtADR1-L1 WT (Figure 4D and 4E) and DV (Figure 4F). The combination of r3m and r4m abolished the phospholipid binding of AtADR1-L1 CC^R^ *in vitro*, and 2 or 5 times more r3m/r4m protein could not restore the binding to WT level (Figure 4A). Moreover, r3m/r4m abolished the cell death activity of AtADR1-L1 DV (Figure 4B and 4C) and dramatically reduced the PM localization of both AtADR1-L1 WT (Figure 4D and 4E) and DV (Figure 4F, S3C and S3D) to be largely cytoplasmic. Based on an AlphaFold structure model of AtADR1-L1 CC^R^ in the resting state, the two clusters of positively charged residues involved in r3m and r4m are spatially close to each other (Figure S4A). Hence, the α2 and α4 helices collectively contribute to phospholipid binding and PM localization of ADR1 before and after activation.

**Figure 4.**
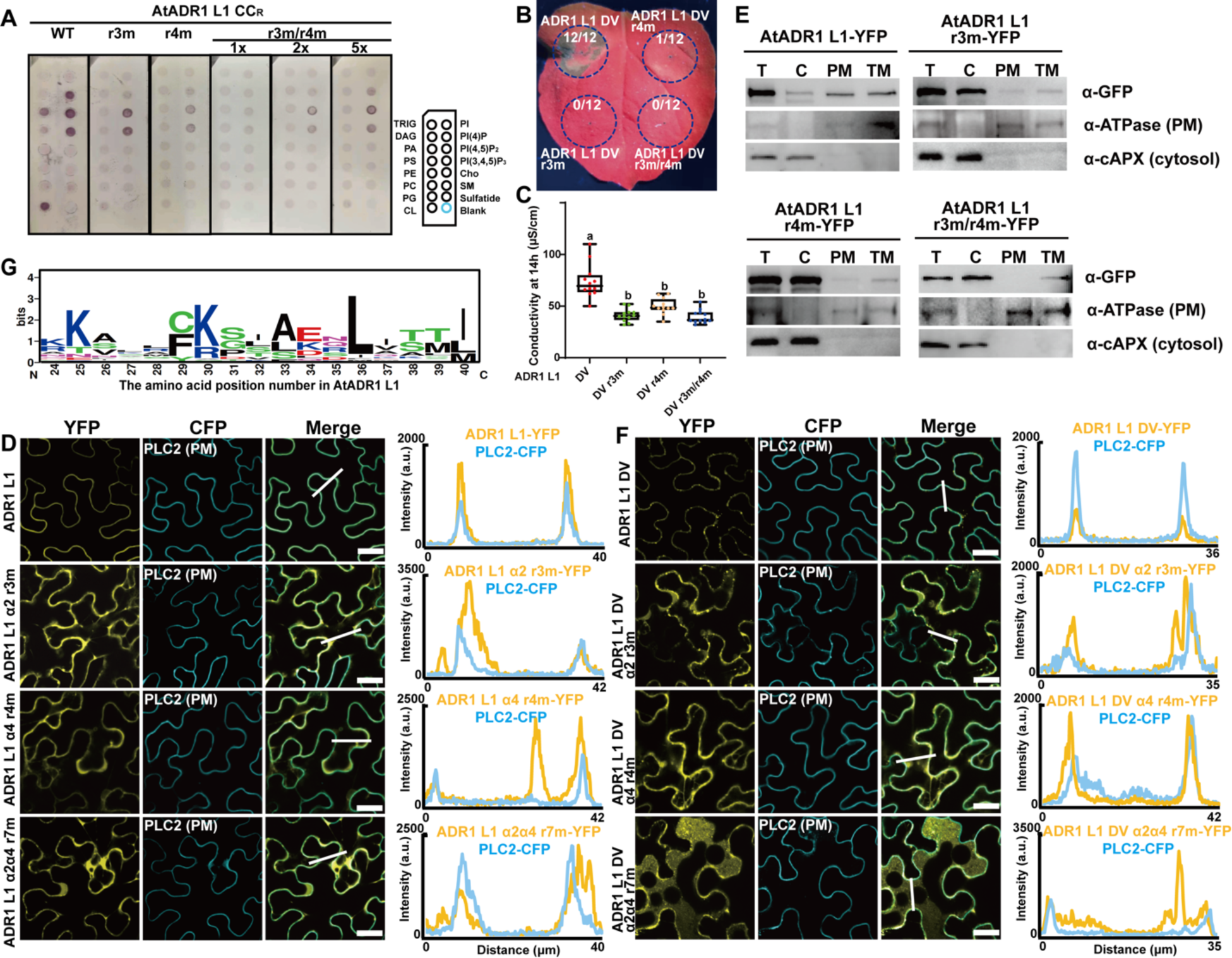
Positively charged residues in the α2 an α4 helices of the 4HB collectively contribute to AtADR1-L1 phospholipid binding, PM localization and function. **(A)** Lipid strip binding assay using purified proteins of AtADR1-L1 CC^R^ (aa 1-114) and mutant derivatives r3m and r4m. See *Methods* for lipid definitions. **(B)** Cell death phenotypes of AtADR1-L1 DV and mutants with altered PM localization in *Nb* 32h post infiltration. (**C**) Quantification of ion leakage from the cell death phenotypes in (B). Confocal microscopy analyses of the mutations of the positively charged residues affecting PM localizations of AtADR1-L1 **(D)** and AtADR1-L1 DV **(F)**, respectively. The indicated proteins fused with a C-terminal YFP were transiently co-expressed with the PM marker PLC2 fused to CFP in *Nb* leaves and confocal images were taken at 32-36h post infiltration. Confocal images are single plane secant views. Merge means merged image between YFP and CFP images. Fluorescence intensities were measured along the white line depicted in the merge images. Bars, 25 µm. **(E)** Membrane fractionation assays on the positively charged residues affecting AtADR1-L1 protein accumulation at the PM. **(G)** Sequence logos showing the conservation of AtADR1-L1 K30 in RNLs from different plant species. The alignment was performed with ClustalX and sequence logos were generated using WEBLOGO.

Our data indicate that these helper NLRs are localized to the PM in the resting state via the positively charged residues in the α2 and α4 helices of the 4HB that interact with phospholipids. According to the AtZAR1 activation model, the α1 helix flips out of the 4HB and inserts into the PM upon activation (13) (Figure S4B). The conserved positively charged residue in the α2 helix corresponding to K30 in AtADR1-L1 is located at the very beginning of α2 (Figure S2A) and could still associate with the PM following an AtZAR1-like activation model (Figure S4B). Also, the α4 helix becomes disordered upon oligomerization but maintains contact with the PM (13) (Figure S4B). These conformational changes are likely to allow positively charged amino acids in the α2 and α4 helices to maintain association with the PM post activation. Hence, helper NLRs rely on the conserved positively charged residues in both the α2 and α4 helices of the 4HB for phospholipid binding and PM localization before and after activation, despite the conformational changes generated by activation.

### Effector activation of a TNL induces AtNRG1.1 resistosome formation at the PM

Our previous study on auto active AtNRG1.1 DV used different loss-of-function mutations, including L134E that abolished oligomerization, ΔN16 that enhanced oligomerization and D3N/E14Q that specifically blocked Ca^2+^ channel activity (3). It remained unknown whether AtNRG1.1 indeed oligomerizes at the PM following activation by an effector and its corresponding TNL. We transiently expressed AtEDS1/AtSAG101/AtNRG1.1 in *Nb epss* (*eds1a, pad4, sag101a, sag101b*) leaves (6) together with the effector XopQ to reconstitute the TNL Roq1 cell death phenotype (34). AtNRG1.1 WT, but not L134E, ΔN16, D3N/E14Q or g4m, caused cell death (Figure 5A, 5B and S5A). We employed Blue-Native PAGE to investigate AtNRG1.1 oligomerization status following effector activation. AtNRG1.1 WT, ΔN16, D3N/E14Q and g4m oligomerized following Roq1 activation by XopQ, with ΔN16 exhibiting much stronger oligomerization signal than the others; L134E failed to oligomerize (Figure 5C). Hence, g4m specifically affects PM localization of AtNRG1.1, but not oligomerization (Figure 1F, S2 and 2E). These cell death phenotypes and oligomerization states are consistent with the observations using the activation mimic allele AtNRG1.1 DV (Figure 1C, 1D, and 1F) (3). In confocal assays, AtNRG1.1 WT, D3N/E14Q and g4m, which all maintained the ability to oligomerize, also exhibited puncta close to or on the PM, and ΔN16 displayed enhanced puncta; the loss-of-oligomerization mutant L134E exhibited no obvious puncta formation compared to AtNRG1.1 WT (Figure 5D-5I and S5B). These data demonstrate that effector activation of a TNL induces AtNRG1.1 resistosome formation at the PM and that AtNRG1.1 DV faithfully mimics the functionality of effector activated AtNRG1.1.

**Fig. 5.**
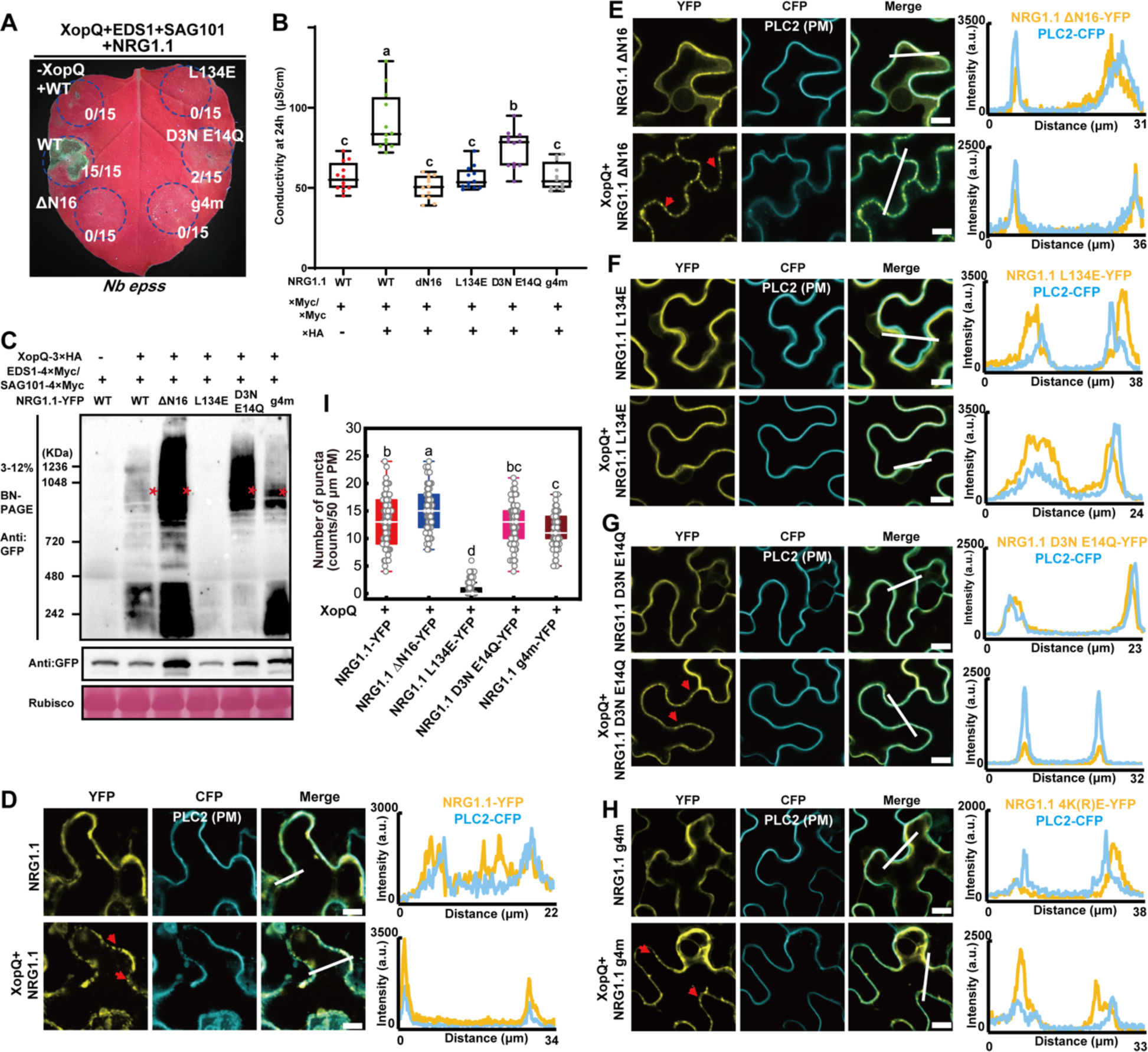
Effector induced AtNRG1.1 oligomerization and enhanced puncta formation at the PM. **(A)** XopQ-triggered cell death phenotypes in *Nb epss* when co-expressed with AtEDS1, AtSAG101 and AtNRG1.1 mutants 48h post infiltration. (**B**) Quantification of ion leakage from the cell death phenotypes in (A). **(C)** BN-PAGE showing the oligomerization status of AtNRG1.1 mutants in (A). Red asterisks indicate oligomerized AtNRG1.1. Confocal microscopy assays showing PM localization and puncta of AtNRG1.1 **(D)**, derivative alleles ΔN16 **(E)**, L134E **(F)**, D3N/E14Q **(G)** and g4m **(H)** before and post-activation via XopQ expression when co-expressed with AtEDS1 and AtSAG101. The indicated proteins fused with a C-terminal YFP were transiently co-expressed with the PM marker PLC2 fused to CFP in *Nb* leaves and confocal images were taken at 32-36h post infiltration. Confocal images are single plane secant views. Images are single plane secant views. Merge means merged images between YFP and CFP images. The red arrow heads indicated puncta. Fluorescence intensities were measured along the white line depicted in the merge images. Bar, 10 µm. (**I**) Quantification of puncta as observed in (D-H).

### The cytoplasmic pools of AtEDS1 and AtSAG101 mediate cell death but are not detectable in the oligomerized AtNRG1.1 resistosome

AtEDS1 and AtSAG101 are localized to the nucleus and the cytoplasm (35, 36). We investigated whether enforced nuclear and cytoplasmic distribution of AtEDS1 and AtSAG101 could affect the cell death phenotype using co-expression complementation assays in *Nb epss* plants. Expression of AtEDS1 and AtSAG101 with a nuclear export signal (NES) slightly attenuated XopQ-TNL Roq1 dependent cell death comparable to wild type AtEDS1 and AtSAG101 (Figure 6A, 6B, and S6), which agrees to some extent with previous finding showing that co-expression of NES-tagged Solanaceae EDS1 with Solanaceae SAG101 blocks XopQ trigged cell death function in *Nb epss* (37). By contrast, AtEDS1 and AtSAG101 with a nuclear localization signal (NLS) did not support cell death in this assay (Figure 6A, 6B, and S6). These data suggest that the cytoplasmic pools of AtEDS1 and AtSAG101 activate AtNRG1.1 and are critical determinants of XopQ-activated cell death responses, which is consistent with a recent study showing that different TIR domains preferentially signal cell death via the cytosolic pool of EDS1 in Arabidopsis (38). Given that TIR-dependent signals induce the formation of AtEDS1/AtSAG101/AtNRG1 heterotrimers (20), we performed BN-PAGE to further investigate if AtEDS1 and AtSAG101 are subsequently retained in the oligomerized AtNRG1.1 resistosome. AtNRG1.1 ΔN16 oligomerized in a XopQ-dependent manner following activation of the Roq1 TNL when co-expressed with AtEDS1 and AtSAG101, while AtNRG1.1 ΔN16/L134E was not responsive to XopQ (Figure 6C and S7). However, in samples expressing AtNRG1.1 ΔN16, with or without XopQ, or AtNRG1.1 ΔN16/L134E plus XopQ, we did not detect oligomerization of AtEDS1 (Figure 6C, StrepII immunoblot) or AtSAG101 (Figure 6C, Flag immunoblot) at the high molecular weight regions corresponding to AtNRG1.1 oligomers. We ruled out the tag effect for the negative results by detecting oligomerization of AtRNG1.1 ΔN16 with the same epitope tag as either AtEDS1 or AtSAG101 in a single gel at three different time points 26h, 32h, and 42h post infiltration (Figure 6C and S7). Hence AtEDS1 and AtSAG101 could not be detected in the oligomerized AtNRG1.1 resistosome in this cell death reconstitution assay in *Nb epss*. One hypothesis is that additional signals may be required to trigger the dissociation of AtEDS1/AtSAG101 and subsequent oligomerization of AtNRG1.1 after TIR-induced AtEDS1/AtSAG101/AtNRG1.1 heterotrimer formation (Figure 6D).

**Fig. 6.**
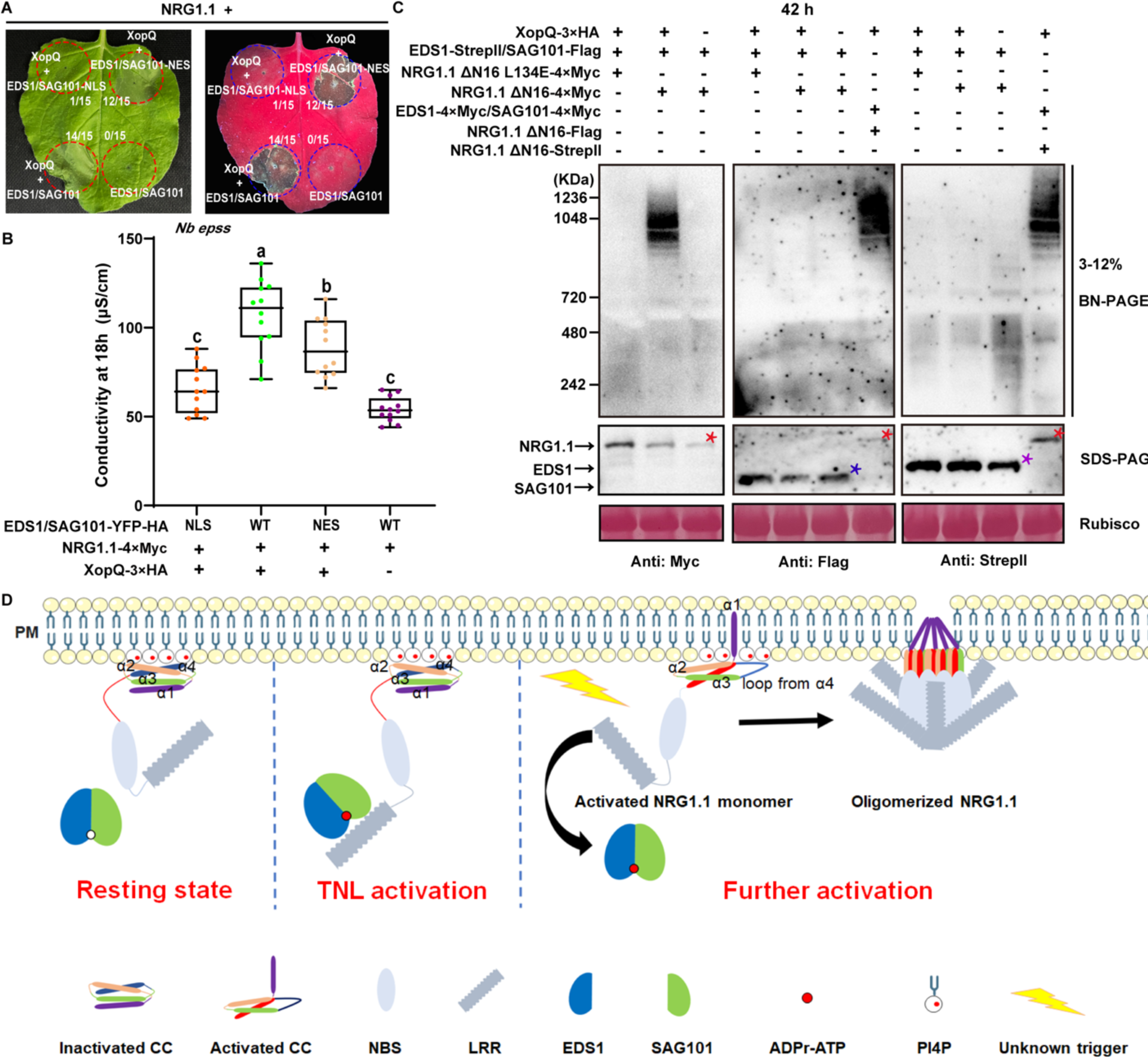
AtEDS1 and AtSAG101 cannot be detected in the oligomerized AtNRG1.1 resistosome. **(A)** The cytoplasmic pool of EDS1/SAG101 activates NRG1.1 to induce cell death. EDS1/SAG101-NLS, EDS1/SAG101-NES, or EDS1/SAG101 with a C-terminal YFP-HA tag were transiently co-expressed with NRG1.1-4×Myc and XopQ-4×Myc in *Nb epss*. Cell death phenotypes were photographed 42h post infiltration. (**B**) Quantification of ion leakage from the cell death phenotypes in (A). **(C)** EDS1/SAG101 do not co-migrate with NRG1.1 upon XopQ activation in BN-PAGE assays. *Agrobacteria* mixes carrying the indicated expression constructs were co-infiltrated into *Nb epss*. Total protein extracts were resolved by BN-PAGE and SDS-PAGE 42h post infiltration. Target bands were immunoblotted with appropriate antibodies. Red asterisk indicates the protein band of NRG1.1. Purple and blue asterisks indicate EDS1 and SAG101 bands, respectively. **(D)** An updated model on PM association and effector-induced oligomerization of NRG1.

## Discussion

PM localization was required for Ca^2+^ channel formation or immune function of RNLs and many CNLs (3, 12, 14, 15, 25, 26, 29, 30). Helper NLRs, including NRG1s and ADR1s, exhibited significant PM location before activation (or the extra-haustorial membrane in the case of NbNRC4) and are further enriched and concentrated at the PM after activation (3, 29-31). Mechanisms driving helper NLR PM localization remained largely unclear.

In plants, PI4P is the main driver of PM electrostatics (39, 40). Phosphatidic acid (PA) and phosphatidylserine and PI(4,5)P_2_ also contribute to PM (inner leaflet) surface charges, but to a lesser extent. We generated mutations in the conserved positively charged residues in the α2 and α4 helices of helper NLR 4HBs. These mutations attenuated phospholipid binding *in vitro* and PM association *in vivo*; they also abolished cell death function. Another 4HB-containing membrane pore forming protein, the animal mixed lineage kinase domain-like (MLKL), specifically interacts with phospholipids for PM localization and function (41). A recent study showed that *Mycobacterium tuberculosis* inhibits pyroptosis by secreting a phospholipid phosphatase that localizes at the host PM to dephosphorylate PI4P and PI(4,5)P_2_ and suppresses the PM association and pore function of activated Gasdermin D (42). Hence phospholipid-binding mediated PM association is a conserved mechanism for membrane pore forming proteins. It remains to be investigated whether plant pathogens employ similar phospholipid phosphatases to perturb the PM localization and thus the function of helper NLRs.

We provide mechanistic insights into the dynamics of PM association during plant helper NLR activation. In the absence of pathogen infection, interactions of the α2 and α4 helices of the N-terminal 4HB anchor a significant fraction of plant helper NLRs at the PM. Upon activation by senor NLRs, helper NLRs undergo dramatical conformational changes and further concentration at the PM; they still rely on the α2 and α4 helices to associate with the PM for function (Figure 5C).

The RNL helper ADR1s and NRG1s contain an RPW8-like CC^R^ domain at their N-termini. In addition to the 4HB, the CC^R^ domain has an extra N-terminal region that was previously predicted to be responsible for CC^R^ membrane targeting (43). Deletion of the extra N-terminal region did not affect PM localization of the activation mimic allele AtNRG1.1 DV (3) or effector-activated AtNRG1.1 (Figure 4D). Mutation of the positively charged residues in the α4 helix affected PM localization of AtNRG1.1 WT and the DV allele (Figure 1D and 1E). Despite the absolute conservation of AtNRG1.1 K100 in the CC^R^ domain of ADR1s and NRG1s, it is not conserved in RPW8 (Figure S1A), indicating a different PM association mechanism for RPW8.

AtNRG1.1 K100 is highly conserved in RNL and NRC helpers, but is less well conserved in sensor CNLs (Figure S1A). In contrast to RNL helpers, the well-studied sensor CNL AtZAR1 that also functions as a PM-localized Ca^2+^ channel post activation, localizes mainly in the cytoplasm before activation (Figure S1B) (12, 13). In contrast, the sensor CNL AtRPM1 interacts with the PM-localized guardee RIN4, localizes to and signals from the PM (25, 44, 45); the senor CNL AtRPS5 localizes to the PM via N-terminal acylation signal and functions at the PM (26). Thus, sensor NLRs express variable resting state subcellular distributions, likely reflecting the subcellular localization of the pathogen effectors they detect. Hence, helper NLR use a specific mechanism for enrichment at the PM before activation, which may be critical for rapid calcium signal induction upon activation of ETI. It remains to be determined whether other CNLs also use similar phospholipid binding mechanisms to determine or enhance PM localization.

PTI and ETI function cooperatively and potentiate each other (46, 47), but the mechanisms that interconnect PTI and ETI remain unclear. The EDS1/PAD4/ADR1 module associates with some PM located PRRs and is required for PTI (48-50). NbNRC3 mediates the hypersensitive cell death caused by the cell-surface receptor Cf-4 recognizing the apoplastic fungal effector Avr4 (51). An enriched or constitutive PM-bound pool of helper NLRs are potentially important for their efficient activation by PTI signals and the cooperative function between PTI and ETI. We demonstrated that NbNRCX, a negative regulator of NbNRC2 and NbNRC3 (32), also uses the conserved positively charged residues in the α4 helix of 4HB for PM association (Figure 2D). We propose that the PM location of NbNRCX is important for its negative regulation of activated NbNRC2 and NbNRC3 at the PM, perhaps via the formation of mixed and thus ‘poisoned’ oligomers.

Effector activation of sensor CNL triggers helper NRC oligomerization, and sensor CNLs are not present in the NRC oligomer complex (52, 53). We present data that ETI activated via the effector XopQ and the TNL Roq1 induces RNL AtNRG1.1 oligomerization and puncta formation at the PM (Figure 4). A recent study showed that in Arabidopsis the oligomerization of AtNRG1.2 requires not only effector-mediated activation of a TNL, but also activation of PTI (54). Our data show that the cytoplasmic pool of AtEDS1/AtSAG101 mediates the cell death response, but that neither are readily detectable in the effector-induced oligomerized AtNRG1.1 resistosome (Figure 5A and 5B). This is in contrast to the recent observation of NRG1.2/EDS1/SAG101 hetero-oligomers that were detectable only after immunoprecipitation of NRG1.2 (54). The combined data suggest that additional unknown triggers may be required to induce AtEDS1/AtSAG101 dissociation from the AtNRG1.1/AtEDS1/AtSAG101 heterotrimer and subsequent AtNRG1.1 oligomerization at the PM (Figure 6D), or that the conformational changes resulting from oligomerization abrogate the interface(s) required for AtEDS1/AtSAG101 co-association. It is also possible that NRG1/EDS1/SAG101 hetero-oligomers are transient, while we detected a latter stage of the oligomerized NRG1 resistosome after EDS1/SAG101 disassociate. The putative role of PTI in facilitating either the rare NRG1.2/EDS1/SAG101 hetero-oligomers or oligomeric NRG1 resistosome formation at the PM requires future investigation.

## Materials and methods

### Plant growth conditions

*Nb* WT and *Nb epss* plants were grown in a growth chamber at 25°C under a 16 h/8 h light/dark cycle with relative humidity at 55±10%.

### Vector construction

The coding sequences (CDS) of *AtNRG1.1* (At5g66900), *AtEDS1* (At3g48090), *AtSAG101* (At5g14930), *AtADR1-L1* (At4g33300) and *PLC2* (At3g08510) were amplified using specific primers from the Arabidopsis ecotype Col-0, and the CDS of *NbNRC4* (NbS00016103g0004, 2,464 bp) and *NbNRCX* (NbS00030243g0001, 2,619 bp) were amplified by PCR from *Nb*. The resulting PCR fragments were cloned into a modified pUC19 vector with the gateway compatible recombination sites (attL1/attL2). The StrepII tag sequence was fused to the 3’ cDNA of *AtEDS1* to obtain an *AtEDS1-StrepII* entry clone. *AtNRG1.1Δ16* was PCR-amplified using the CDS of *AtNRG1.1* as a template and then cloned into the entry vector pUC19. Other *AtNRG1.1* variants were generated using Fast Mutagenesis System protocol (Transgene). *XopQ* with *attL1/attL2* sequences was synthesized into a pUC18 plasmid (Genescript). *XopQ* was introduced to expression vectors pGWB614 and pGWB617 to obtain *35S:: XopQ-3×HA* and *35S:: XopQ-4×Myc* constructs, respectively. *AtEDS1* and *AtEDS1-StrepII* were recombined into pGWB617 (35S promoter, C-terminal 3×Myc) and pGWB2 (35S promoter, no tag), respectively. *AtSAG101* was introduced to pGWB11 (35S promoter, C-terminal Flag) and pGWB617. *AtNRG1.1* and the mutants were introduced into pGWB641 (35S promoter, C-terminal EYFP), pGWB617, the modified vectors pEarleyGate-Flag (35S promoter, C-terminal Flag) and pEarleyGate-StrepII (35S promoter, C-terminal StrepII) to express fusion proteins with desired tags. All target sequences were introduced into expression vectors using LR reactions (Life Technologies). *35S::AtEDS1-YFP-HA-NLS* (or *NES*) and *35S::AtSAG101-YFP-HA-NLS* (or *NES*) expression constructs were obtained using LR to recombine pUC19-*AtEDS1*, pUC19-*AtSAG101* with modified vectors pEarleyGate 101-NLS and pEarleyGate 101-NES (55). *NbNRC4* and *NbNRCX*, and their mutants were introduced into the 35S-omega-driven destination vector pEarlyGate with a C-terminal YFP-HA. The CDS of plasma membrane marker PLC2 (At3g08510) and endoplasmic reticulum marker VMA12 (At5g52980) were amplified from Col-0 and subsequently cloned into PUC19/Kan vector including attR1/attR2 sites. *PLC2-CFP* expression vector and *VMA12-RFP* were generated by Golden Gate assembly with 35S-driven pEarly101 with a C-terminal CFP and 35S-driven pGWB257 with a C-terminal RFP, respectively. The obtained expression constructs were transformed into *Agrobacterium tumefaciens* GV3101 via electroporation for transient expression in *Nb*.

### Transient expression in *Nb*

*Agrobacteria* (GV3101) carrying the constructs were grown overnight in LB with suitable antibiotics. 3 mL of *Agrobacterium* culture was centrifuged and resuspended in MES buffer (10 mM MgCl_2_, 10 mM MES pH 5.6, 150 μM acetosyringone). *Agrobacteria* were incubated at room temperature for 1h and infiltrated into the leaves of 5-week-old *Nb or Nb epss* at specific OD_600_ values. *Agrobacteria* containing the *P19* construct was co-infiltrated at OD of 0.1. Cell death phenotypes were photographed at indicated time point.

### Ion leakage assays

Four leaf discs 8mm in diameter were harvested 24h after transient expression in *Nb* leaves. Leaf discs were washed for 1h in 10 mL ultrapure water, transferred into 15-mL tubes containing 6 mL ultrapure water. Conductivity was measured using a conductivity meter (FiveGo Cond meter F3, METTLER TOLEDO) at indicated time point. 3 biological repeats and 4 technical repeats were performed. Statistical analysis was performed via Tukey’s HSD (honestly significant difference) using GraphPad Prism 8. Different letters indicate statistically significant differences.

### Total protein extraction and western blot analysis

Three 8 mm leaf discs were harvested and ground to powder in liquid nitrogen. Total protein was extracted with 100 μL extraction buffer (20 mM Tris–HCl pH 8.0, 5 mM EDTA, 1% SDS, 10 mM DTT). Lysate was boiled at 95 °C with 1×protein loading buffer for 10 min. The total protein extract was cleared by centrifuge at 13,000 *g* for 10 min. Then the supernatant was separated by 10% SDS–PAGE gels and detected with corresponding antibodies. Antibodies used for immunoblotting include anti-GFP (#G1544, Sigma), anti-HA (#11867423001, Roche), anti-Myc (#M4439, Sigma), anti-Flag (#SAB4301135, Sigma), anti-StrepII (#ab76949, Abcam), and HRP-conjugated antibodies (#IMR-GtxMu-003-DHRPX, #IMR-GtxRb-003-DHRPX, #GtxRt-003-DHRPX, Jackson).

### BN-PAGE

BN-PAGE was performed as previously described with slight modifications (3). 3 leaf disks (5 mm diameter) were homogenized with liquid nitrogen. 1× NativePAGE™ Sample Buffer (BN2008, Invitrogen™) with 1× protease inhibitor cocktail was added to the homogenized samples. The mixed samples were centrifuged at 20,000 *g* for 30 min at 4°C. Native PAGE™ 5% G-250 Sample Additive (BN2004, Invitrogen™) was added to the supernatant at a final concentration of 0.125%. Proteins were separated by Native PAGE™ Novex® 3–12% Bis-Tris Gels (BN1001, Invitrogen™) and transferred to PVDF membrane using an eBlot™ L1 transfer system (GenScript). The target proteins were probed with corresponding antibodies.

### Confocal microscopy analyses

For *Agrobacterium*-mediated transient expression in *Nb*, OD_600_ cultures were adjusted to 0.1 of P19, 0.2 of PLC2, and 0.2 of AtNRG1.1, AtADR1-L1, NbNRC4 or NbNRCX. *Agrobacteria* mixtures were infiltrated into young leaves of 4-6 weeks old *Nb* plants. For transient expression in *Nb epss*, the OD_600_ was adjusted to 0.1 of P19, 0.1 of AtEDS1 and AtSAG101, and 0.2 of AtNRG1.1 and PLC2 (VMA12-RFP or RFP), and 0.001 for XopQ. Agrobacteria mixtures were infiltrated into young leaves of 4-6 weeks old *Nb epss* plants. Leaves were imaged for protein localization analyses between 36h-48h post infiltration. Confocal images were taken on a confocal laser scanning microscope LSM880 from Zeiss (Oberkochen, Germany) using the ZENblack software, via a Zeiss-C-Apochromat 20×/0.8 A20650-9901 objective or a Zeiss-W-Plan-Apochromat 421462-9900 objective. YFP was excited using a 514 nm laser and the emission spectrum was between 516-556 nm; CFP was excited with a 458 nm laser and the emission spectrum was between 463-513 nm. Focal plane images were processed with the ZEN blue software (Zeiss) for adjustment of brightness and contrast.

### Membrane fractionation assays

Plasma membrane protein isolation was carried out by slightly modifying a previously described protocol (3).In brief, sucrose buffer [20mM Tris (pH 8.0), 0.33M sucrose, 1mM EDTA, 5mM DTT and 1x Sigma plant protease inhibitor cocktail] was added to the homogenized tissue at a ratio of 5 uL per mg (FW) tissue. The extract was centrifuged at 2,000 x *g* for 5 minutes at 4°C, then the supernatant was transferred to a new tube and designated as total protein (T). Cytoplasmic fraction (C) was prepared by harvesting the supernatant after spinning the total protein fraction at 20,000 x *g* for 1h at 4°C. The total membrane fraction (TM) was prepared from the resulting pellet by resuspending in 200 uL of buffer B (Minute^TM^ plasma membrane protein isolation kit, Invent Biotechnologies). Centrifuged at 7,800 x *g* for 15 min at 4°C, then the supernatant was transferred to 2 mL Eppendorf tube and mixed with 1.6 mL cold PBS buffer mixed by vortexing and spun at 16,000 x *g* for 1h at 4°C to pellet the plasma membrane fraction. The pellet was resuspended with sucrose buffer in 4 times less volume than the soluble fraction. The resulting fraction was labelled as the plasma membrane-enriched/microsomal fraction (PM). Proteins fractions were run on SDS-PAGE gels and analyzed by western blotting.

### Sequence alignment of helper NLR homologs

AtNRG1.1 and AtNRG1.2 (At5g66910.1) were used to identify NRG1 homologs in the predicted protein databases including Solanaceae Genomics Network (SGN) and Integrated Microbial Genomes (IMG). Similarly, AtADR1 (At1g33560.1), AtADRL1-L1 and AtADRL1-L2 (At5g04720.1 were used to identify ADR1 homologs. NbNRC2 (NbS00018282g0019.1 and NbS00026706g0016.1), NbNRC3 (NbS00011087g0003.1), NbNRC4 (NbS00002971g0007.1 and NbS00016103g0004.1) and NbNRCX (NbS00030243g0001.1) were used to identify NRC-helper clade homologs in the SGN database. Top hits from BLASTP search results in different plant species were collected for further analyses. Homologs with protein sequence identity of more than 50% were aligned in their CC or CC^R^ domains by CLustalW. The gene ID of homologs used in the alignment were listed in Table S1.

### Sequence logos visualization

Sequence logos of the identified conserved amino acid site in CC or CC^R^ domain of helper NLRs were generated by the online software WEBLOGO (http://weblogo.berkeley.edu/logo.cgi) (56).

### Protein expression and Lipid-strip binding assays

Sequences of AtNRG1.1 CC 1-124 (WT, R99E, K100E, R103E, K106E and K110E) were subcloned into the pET24a plasmid with a N-terminal 6×His and a C-terminal StrepII tag. Proteins were expressed in *E. coli* BL21 (DE3) cells using the autoinduction method (57). Cell pellets were resuspended and lysed in buffer containing 50 mM HEPES (pH 8.0), 300 mM NaCl, 2 mM DTT and 1 mM PMSF. After sonication and centrifugation at 20,000 *g* for 60 minutes, supernatant was loaded onto a nickel HisTrap 5 mL column (GE Healthcare) pre-equilibrated with 20 mL of the wash buffer (50 mM HEPES pH 8.0, 300 mM NaCl, 30 mM imidazole) at a 3 mL/min rate. The bound proteins were eluted with elution buffer (50 mM HEPES pH 7.5, 300 mM NaCl, 500 mM imidazole), and further purified using size exclusion chromatography (Superdex 200 HiLoad 16/600 column) pre-equilibrated with the buffer containing 10 mM HEPES pH 7.5, 150 mM NaCl and 1mM DTT. The peak fractions were confirmed by SDS-PAGE and pooled for Lipid-strip binding assays.

Proteins of AtADR1 L1 CC^R^ 1-114 (WT, r3m, r4m and r3m/r4m), NbNRC4 CC 1-127 (WT and c5m) and NbNRCX CC 1-129 (WT and x4m) were expressed and purified in the human embryonic kidney cell line 293F. The corresponding cDNAs of these constructs were subcloned into the pMlink vector with a N-terminal proteinA and SUMOstar tag and a C-terminal Flag tag. Plasmids were transfected into 293F cells using the polyethyleneimine (PEI) method (58). Cells were grown in Union-293 Chemically Defined Medium (Union Bio) and were routinely maintained at exponential phase in 1000-mL shaker flasks. The flasks were agitated at 130 rpm at 37°C in a humidified condition containing 5% CO2. Cells were pelleted at 8000 x g at 4°C for 5 minutes using centrifugation at 72h after transfection. Pellets were resuspended and lysed in lysis buffer (50 mM Tris-Hcl, 300 mM NaCl, 0.5 mM EDTA, 10 % (v/v) Glycerol, 2 mM DTT, 1 mM PMSF, 5 mM ATP, 2 mM MgSO4, 1 % (v/v) Cocktail, 0.9 % (m/v) DNase I and 0.2 % (m/v) CHAPS, pH 7.5). After vibration at 4°C for 60 minutes and centrifugation at 20,000 x g for 60 minutes, supernatant was loaded onto 6 mL immunoaffinity chromatography column with 1 mL IgG beads (Smart-Lifesciences) pre-equilibrated with 50 mL of the wash buffer (50 mM Tris-Hcl, 300 mM NaCl, 0.5 mM EDTA, 10 % (v/v) Glycerol, pH 7.5). The bound proteins were digested by ULPI enzyme, and further the flow-through containing approved proteins were collected and concentrated for Lipid-strip binding assays.

Lipid-strip binding assays were performed according to the manufacturer’s instructions (Echelon Biosciences, Salt Lake City, UT, USA). Briefly, the PIP-strip membranes were blocked overnight at 4 °C in blocking buffer containing 0.1% Tween-20 and 4% fatty acid-free bovine serum albumin in PBS buffer. Purified proteins (0.5 μg/mL, final concentration) were incubated with the PIP-strip membranes for 1 h at room temperature and then washed three times using wash buffer containing 0.1% Tween-20 in PBS buffer. Bound proteins were detected by immunodetection of StrepII. Lipids in the strip include phosphatidylinositol (4)-phosphate (PI(4)P), phosphatidylinositol (4,5)-bisphosphate (PI(4,5)P2), phosphatidylinositol (3,4,5)-trisphosphate (PI(3,4,5)P3), triglyceride (TRIG), diacylglycerol (DAG), phosphatidic acid (PA), phosphatidylserine (PS), phosphatidylethanolamine (PE), phosphatidylcholine (PC), phosphatidylglycerol (PG), cardiolipin (CL), phosphatidylinositol (PI), cholesterol (Cho), sphingomyelin (SM) and 3-sulfogalactosylceramide (Sulfatide).

## Author Contributions

L.W., J.L.D, Z.W., X.L. designed research. Z.W., X.L., J.Y. S.Y., W.C. performed research. Z.W., X.L., L.W., J.L.D, F.E.K., J.Y., N.H.K. analyzed data. L.W. and J.L.D wrote the paper.

## Competing Interest Statement

The authors declare no competing interests.

## Classification

Plant biology/immunology/biochemistry

## Acknowledgments

We thank the confocal microscopy facility at CAS Ecology/Center for Excellence in Molecular Plant Sciences for technical supports. We thank Professor Jane Parker for providing *Nb epss* seeds. L.W. was supported by National Key Laboratory of Plant Molecular Genetics, Institute of Plant Physiology and Ecology/Center for Excellence in Molecular Plant Sciences and Chinese Academy of Sciences Strategic Priority Research Program (Type-B; Project number: XDB27040214). X.L. was supported by a Postdoc Grant (Project number: 2021M693172) from National Natural Science Foundation of China. F.E-K. is supported by the German Research foundation (DFG Grants CRC1101 D09 and EL 734/3-1). J.L.D. is supported by National Science Foundation (Grant IOS-1758400) and HHMI. J.L.D. is a Howard Hughes Medical Institute (HHMI) Investigator. This article is subject to HHMI’s Open Access to Publications policy. HHMI lab heads have previously granted a nonexclusive CC BY 4.0 license to the public and a sublicensable license to HHMI in their research articles. Pursuant to those licenses, the author-accepted manuscript of this article can be made freely available under a CC BY4.0 license immediately upon publication.

## Supplementary figures

**Figure S1.**
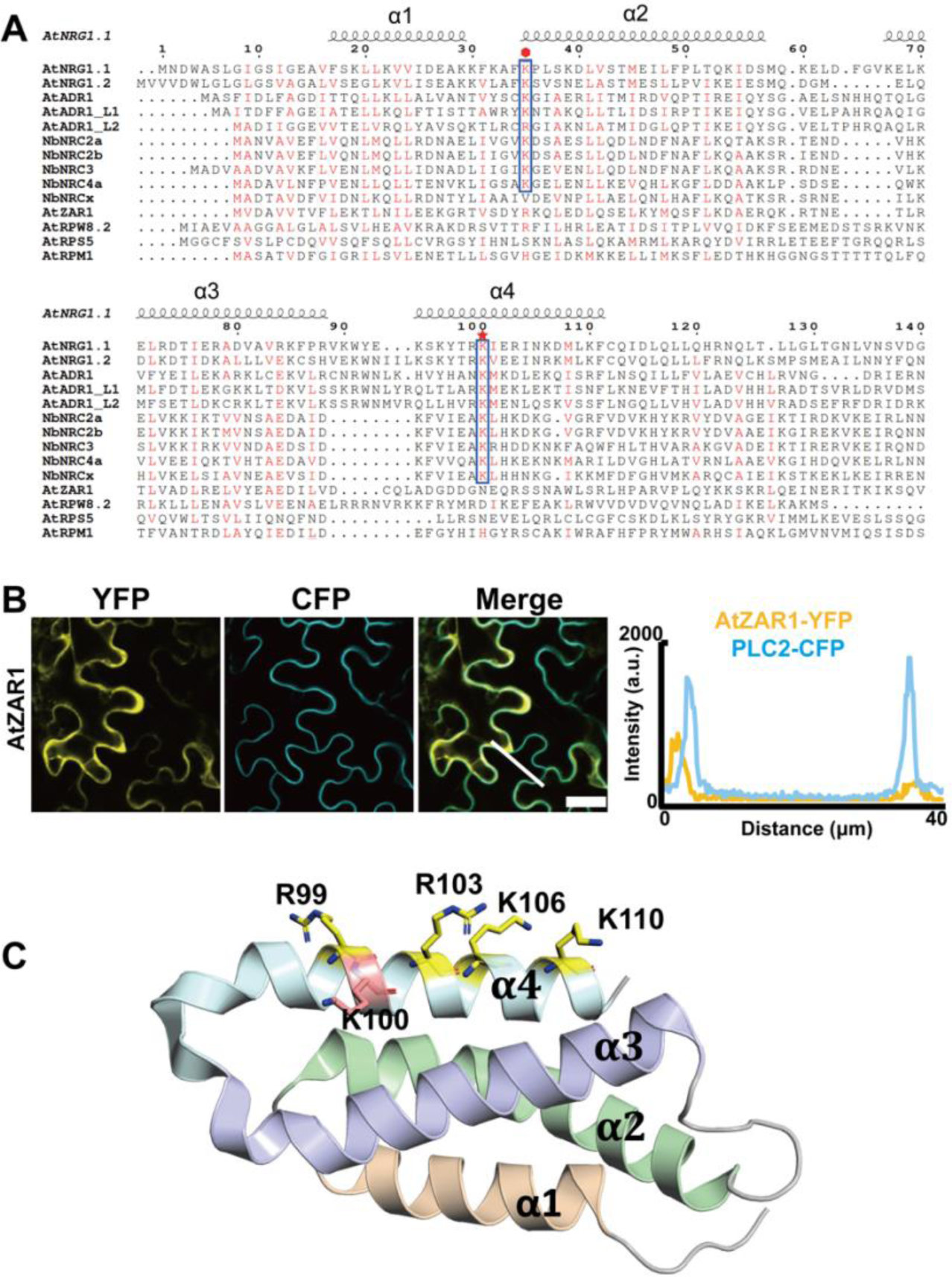
Identification of conserved positively charged residues in plant helper NLRs. **(A)** Sequence alignment in the CC^R^ and CC domains of helper NLRs and some sensor CNLs from Arabidopsis and *Nb*. The alignment was performed with ClustalX and the figure was prepared using ESPript. Red asterisk points to the position of AtNRG1.1 K100, and red hexagon points to the position of AtADR1_L1 K30. **(B)** Confocal assay showing that resting state AtZAR1 mainly localizes in the cytosol. Fluorescence intensities were measured along the white line depicted in the merge images. Bars, 25 µm. **(C)** Cartoon presentation of AtNRG1.1 CC^R^ structure (PDB:7L7W) with R99 K110, R103, K106 and K110 shown as sticks.

**Figure S2.**
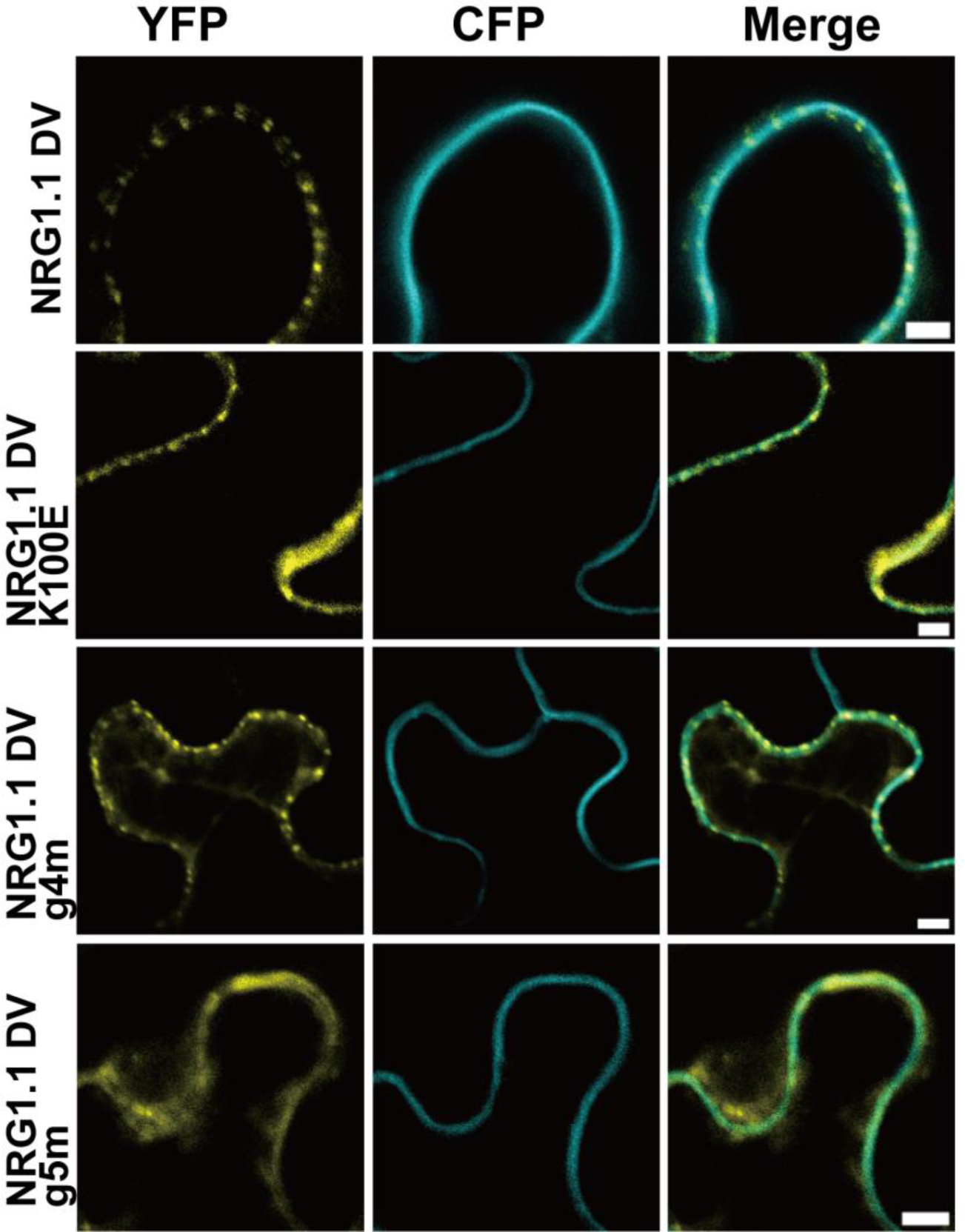
Peripheral images of the confocal assays in Figure 2C.

**Figure S3.**
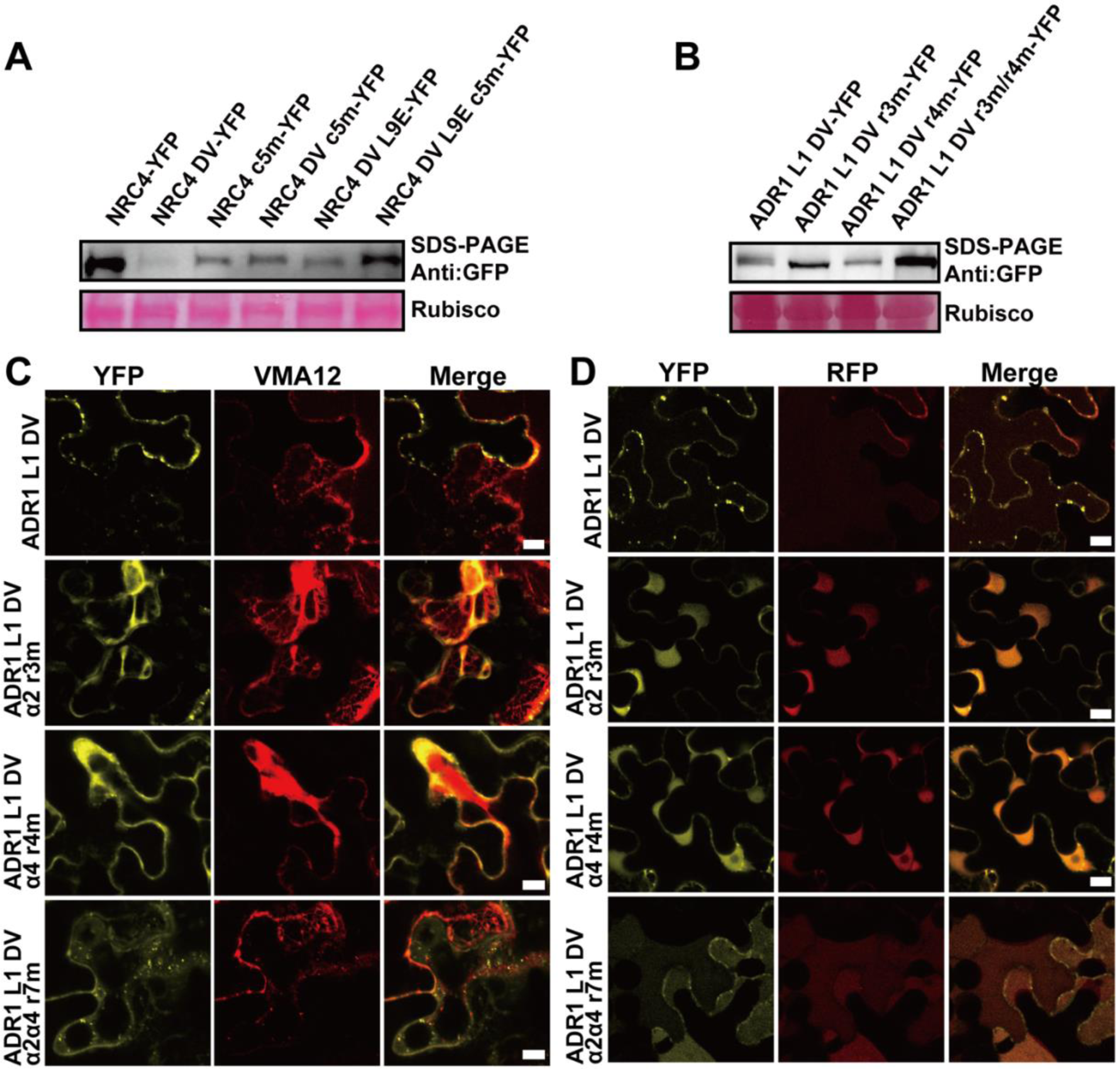
(A) Protein expression levels of NbNRC4 and NbNRC4 DV relevant mutants in Figure 3B. **(B)** Protein expression levels of AtADR1_L1 relevant mutants assayed by immunoblot. Confocal microscopy assays showing if the mis-localized ADR1_L1 DV mutants co-localize with the ER marker VMA12 (**C**) or cytosolic RFP (**D**). The indicated proteins fused with a C-terminal YFP were transiently co-expressed with the ER marker VMA12 fused to RFP or RFP in *Nb* leaves and confocal images were taken at 32-36h post infiltration. Confocal images are single plane secant views. Images are single plane secant views. Merge means merged images between YFP and RFP images.

**Figure S4.**
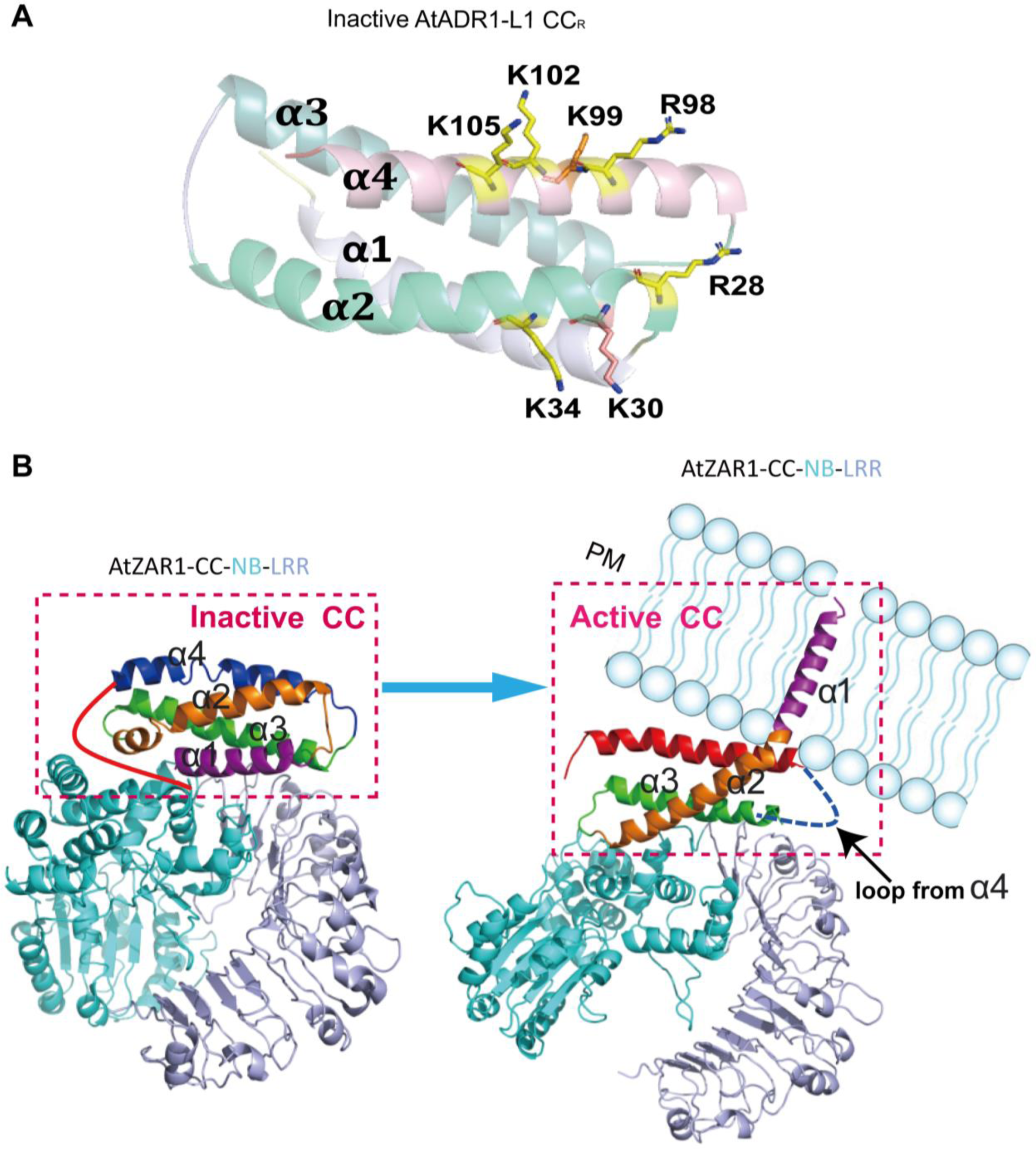
(A) Cartoon presentation of an AlphaFold AtADR1-L1 structure highlighting the positively charged residues involved in r3m and r4m. **(B)** Cartoon presentation of AtZAR1 in resting state (PDB: 6J5W) and in active state (PDB: 6J5T) highlighting the conformational changes in the CC domain and PM association.

**Figure S5.**
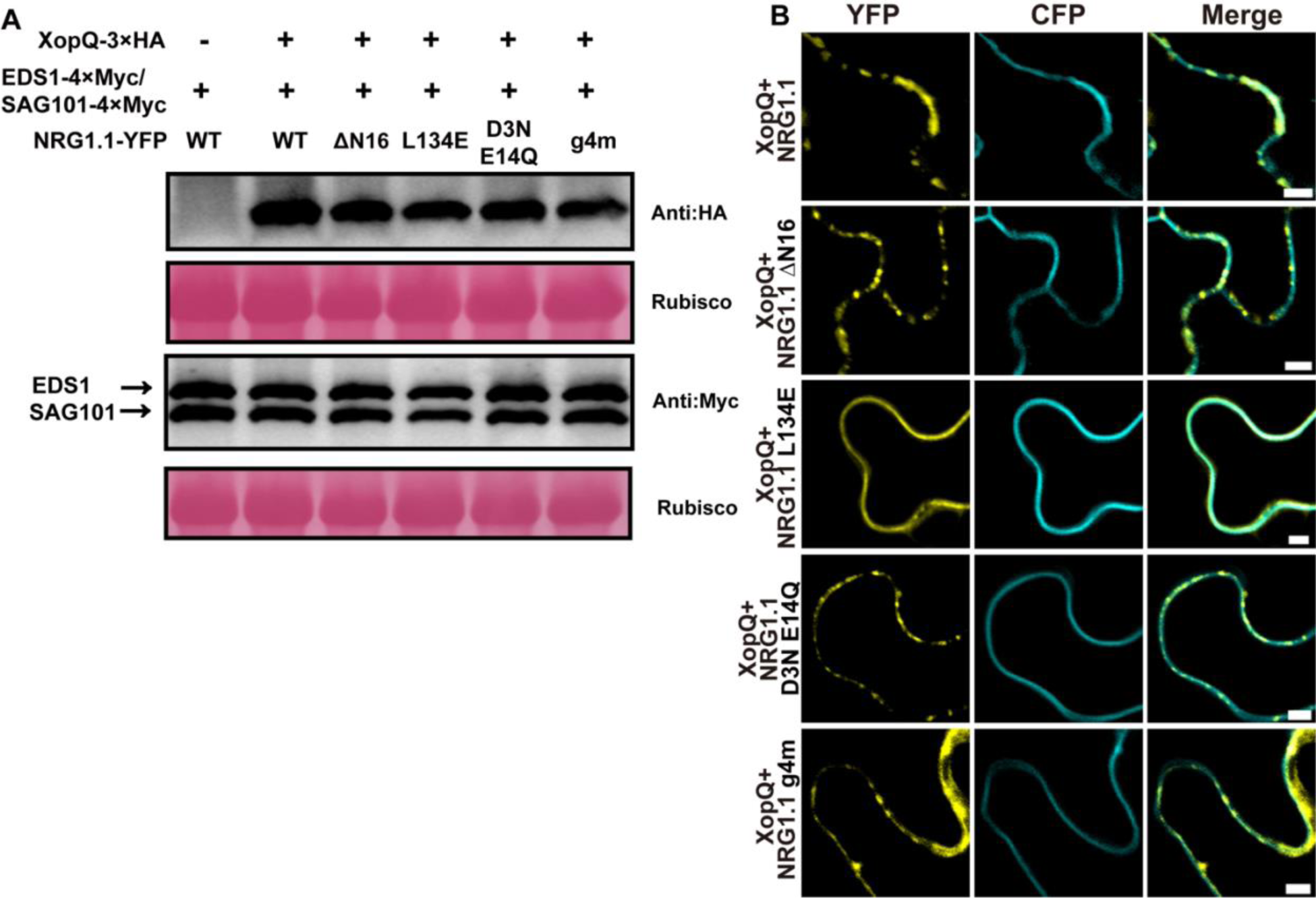
**(A)**Protein levels of XopQ, EDS1 and SAG101 from experiments displayed in Figure 5A and 4B. HA-tagged XopQ and Myc-tagged EDS1/SAG101 were separated by SDS-PAGE and blotted for anti-HA and anti-Myc, respectively. **(B)** Peripheral images of the confocal assays in Figure 5D-5H.

**Figure S6.**
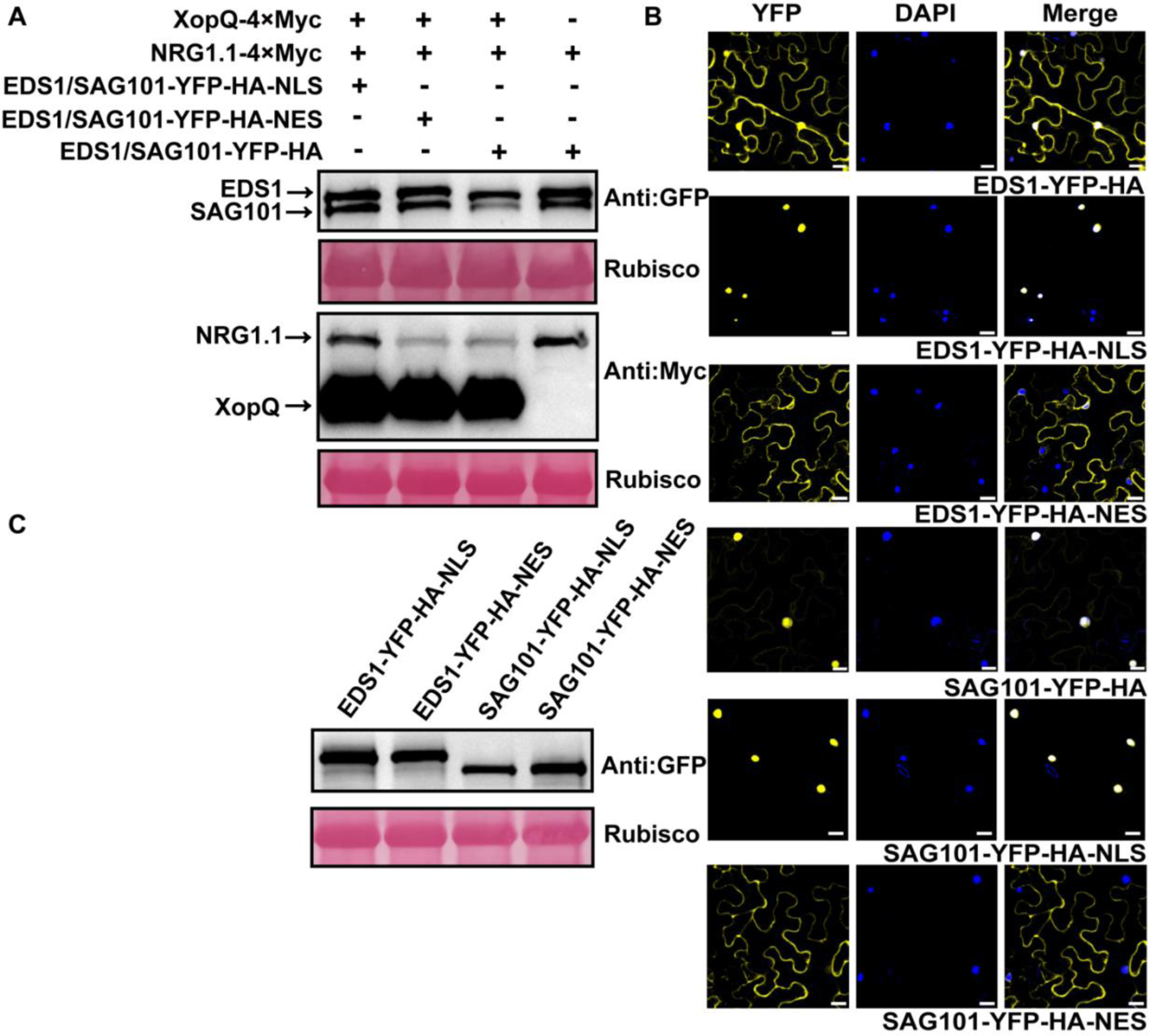
(A) Protein expression levels of samples from experiments displayed in Figure 6A. Total proteins were extracted from samples harvested at 26h post infiltration. Target proteins were separated by SDS-PAGE and blotted with the appropriate antibodies. **(B)** Confocal images showing the subcellular localization of EDS1-YFP-HA, EDS1-YFP-HA-NLS, EDS1-YFP-HA-NES, SAG101-YFP-HA, SAG101-YFP-HA-NLS and SAG101-YFP-HA-NES. Images were photographed at 48h post infiltration in *Nb* leaves. Nuclei were stained with DAPI (4’, 6-diamidino-2-phenylindole). Bars, 20 μm. **(C)** Protein accumulation of samples in (B) detected with anti-GFP antibody.

**Figure S7.**
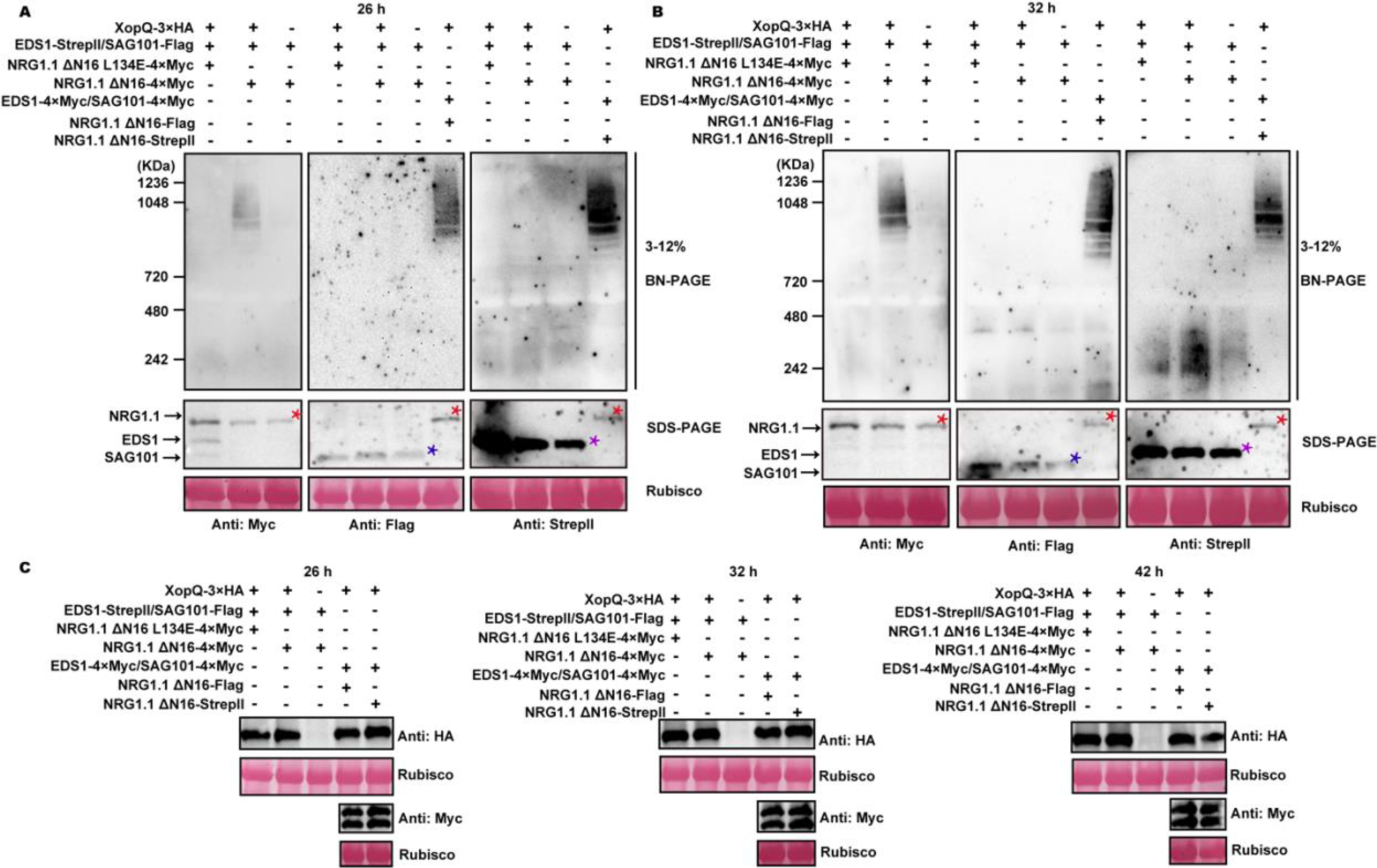
BN-PAGE assays showing that EDS1/SAG101 is not observed in an oligomeric complex with NRG1.1 at 26h **(A)** or 32h (**B)** after co-expression with XopQ in *Nb epss*. Red, purple and blue asterisks indicate NRG1.1, EDS1 and SAG101 bands, respectively. **(C)** Protein expression of XopQ, EDS1 and SAG101 as displayed in Figure 4B and Figure S5A and S5B was verified by SDS-PAGE. XopQ, EDS1/SAG101 were detected with anti-HA antibody and anti-Myc antibody, respectively.

**Table S1.** Gene ID of the 69 helper NLR homologs used in the alignment by CLustalW.

| Gene | Gene ID (databases)<sup>1</sup> | Plant species |
| --- | --- | --- |
| AtNRG1.1 | At5g66900.1 (TAIR) | *Arabidopsis thaliana* |
| AtNRG1.2 | At5g66910.1 (TAIR) | *Arabidopsis thaliana* |
| NbNRG1 | NbS00018358g0013.1 (SGN) | *Nicotiana benthamiana* |
| AINRG1 | 2614272850 (IMG) | *Arabidopsis lyrata* |
| AhNRG1 | 2614358615 (IMG) | *Arabidopsis halleri* |
| CrNRG1 | 2614531086 (IMG) | *Capsella rubella* |
| EhNRG1 | 2615507434 (IMG) | *Eutrema halophilum* |
| CsNRG1 | 2615787498 (IMG) | *Citrus sinensis* |
| CcNRG1 | 2614549921 (IMG) | *Citrus clementina* |
| PpNRG1 | 2615813723 (IMG) | *Prunus persica* |
| VvNRG1.1 | 649552880 (IMG) | *Vitis vinifera* |
| VvNRG1.2 | 649554972 (IMG) | *Vitis vinifera* |
| SpNRG1.1 | 2615336907 (IMG) | *Salix purpurea* |
| SpNRG1.2 | 2615348770 (IMG) | *Salix purpurea* |
| RcNRG1 | 649572805 (IMG) | *Ricinus communis* |
| PtNRG1 | 2615290452 (IMG) | *Populus trichocarpa* |
| SINRG1 | Solyc02g090380.4.1 (SGN) | *Nicotiana benthamiana* |
| AtADR1 | At1g33560.1 (TAIR) | *Arabidopsis thaliana* |
| AtADRL1 L1 | At4g33300.1 (TAIR) | *Arabidopsis thaliana* |
| AtADRL1 L2 | At5g04720.1 (TAIR) | *Arabidopsis thaliana* |
| NbADR1 | Niben101Scf02422g02015.1 (SGN) | *Nicotiana benthamiana* |
| AIADR1 | 2614261905 (IMG) | *Arabidopsis lyrata* |
| AIADR1 L1 | 2614291476 (IMG) | *Arabidopsis lyrata* |
| AIADR1 L2 | 2614268259 (IMG) | *Arabidopsis lyrata* |
| AhADR1 | 2614369556 (IMG) | *Arabidopsis halleri* |
| CrADR1 | 2614512948 (IMG) | *Capsella rubella* |
| CrADR1 L1 | 2614509067 (IMG) | *Capsella rubella* |
| CrADR1 L2 | 2614505477 (IMG) | *Capsella rubella* |
| EhADR1 | 2615515499 (IMG) | *Eutrema halophilum* |
| EhADR1 L1 | 2615525947 (IMG) | *Eutrema halophilum* |
| EhADR1 L2 | 2615515499 (IMG) | *Eutrema halophilum* |
| GrADR1 | 2614796124 (IMG) | *Gossypium raimondii* |
| PtADR1 | 2615278681 (IMG) | *Populus trichocarpa* |
| CcADR1 | 2614532153 (IMG) | *Citrus clementina* |
| CsADR1 | 2615787425 (IMG) | *Citrus sinensis* |
| VvADR1 | 649539454 (IMG) | *Vitis vinifera* |
| SpADR1 | 2615316196 (IMG) | *Salix purpurea* |
| PpADR1 | 2615813687 (IMG) | *Prunus persica* |
| RcADR1 | 649560471 (IMG) | *Ricinus communis* |
| NbNRC2a | NbS00018282g0019.1 (SGN) | *Nicotiana benthamiana* |
| NbNRC2b | NbS00026706g0016.1 (SGN) | *Nicotiana benthamiana* |
| NbNRC3 | NbS00011087g0003.1 (SGN) | *Nicotiana benthamiana* |
| NbNRC4a | NbS00002971g0007.1 (SGN) | *Nicotiana benthamiana* |
| NbNRC4b | NbS00016103g0004.1 (SGN) | *Nicotiana benthamiana* |
| NbNRCX | NbS00030243g0001.1 (SGN) | *Nicotiana benthamiana* |
| SINRC1 | Solyc01g090430.3.1 (SGN) | *Solanum lycopersicum* |
| SINRC2 | Solyc10g047320.2.1 (SGN) | *Solanum lycopersicum* |
| SINRC3 | Solyc05g009630.3.1 (SGN) | *Solanum lycopersicum* |
| SINRC4a | Solyc04g007070.3.1 (SGN) | *Solanum lycopersicum* |
| SINRC4b | Solyc04g007060.3.1 (SGN) | *Solanum lycopersicum* |
| SINRC4c | Solyc04g007030.3.1 (SGN) | *Solanum lycopersicum* |
| SINRCX | Solyc03g005660.4.1 (SGN) | *Solanum lycopersicum* |
| StNRC1a | PGSC0003DMT400066999 (SGN) | *Solanum tuberosum* |
| StNRC1b | PGSC0003DMT400066996 (SGN) | *Solanum tuberosum* |
| StNRC1c | PGSC0003DMT400066993 (SGN) | *Solanum tuberosum* |
| StNRC2a | PGSC0003DMT400080649 (SGN) | *Solanum tuberosum* |
| StNRC2b | PGSC0003DMT400080647 (SGN) | *Solanum tuberosum* |
| StNRC4a_1 | PGSC0003DMT400041024 (SGN) | *Solanum tuberosum* |
| StNRC4a_2 | PGSC0003DMT400069010 (SGN) | *Solanum tuberosum* |
| StNRC4a_3 | PGSC0003DMT400069014 (SGN) | *Solanum tuberosum* |
| StNRC4a_4 | PGSC0003DMT400041023 (SGN) | *Solanum tuberosum* |
| StNRC4b_1 | PGSC0003DMT400019304 (SGN) | *Solanum tuberosum* |
| StNRC4b_2 | PGSC0003DMT400019303 (SGN) | *Solanum tuberosum* |
| StNRC4b_3 | PGSC0003DMT400019302 (SGN) | *Solanum tuberosum* |
| StNRC4b_4 | PGSC0003DMT400019327 (SGN) | *Solanum tuberosum* |
| SmNRC1 | SMEL4.1_12g003240.1 (SGN) | *Solanum melongena* |
| SmNRC3 | SMEL4.1_10g001370.1 (SGN) | *Solanum melongena* |
| SmNRC4a | SMEL4.1_11g018090.1 (SGN) | *Solanum melongena* |
| SmNRC4b | SMEL4.1_11g018170.1 (SGN) | *Solanum melongena* |

<sup>1</sup>TAIR, IMG and SGN are abbreviated from Arabidopsis Information Resource, Integrated Microbial Genomes and Solanaceae Genomics Network databases.

## Notes

### Competing Interest Statement

The authors have declared no competing interest.

### Summary of Updates

Firstly， now quality confocal images have been uploaded with puncta quantified. Secondly, we generated in vitro lipid binding data for NRC4, NRCX and ADR1 (CC domain and mutants). We developed a semi-quantitative assay for lipid binding, in which we showed that higher concentrations of mutant proteins could not restore the binding of mutants to match wild-type proteins of NRG1 and ADR1 CCR. Thirdly, we generated membrane fractionation data on NRG1, NRC4, NRCX and ADR1 to support the confocal assays. Fourthly, we quantified all the HR assays using electrolyte leakage. Finally, we toned down the conclusion of EDS1/SAG101 are not part of the activated complex by saying that EDS1/SAG101 cannot be detected in the effector-activated NRG1 resistosome.

